# Phylogenomics and Systematics of African *Sesuvium* (Aizoaceae)

**DOI:** 10.1101/2025.08.11.669638

**Authors:** Gudrun Kadereit, Alina Höwener, Diana Garaeva, Diego F. Morales-Briones

## Abstract

*Sesuvium* (Sesuvioideae-Aizoaceae) comprises leaf succulent annual and perennial herbs distributed in coastal or saline sites of subtropical and tropical regions. Some species of the genus tolerate highly salinized or polluted soils and show soil-improving properties. *Sesuvium* shows high flexibility of photosynthetic types, including and integrating C_3_, C_4,_ and CAM photosynthesis. Previous molecular phylogenetic studies failed to resolve the infrageneric relationships of the genus. However, a better understanding of the systematics of this group is a prerequisite for further evolutionary studies.

We explored the suitability of genome skimming, including mainly herbarium material, to produce a robust phylogeny of the presumably young African clade of *Sesuvium*. The approach generated an average of 6,289 orthologs per genome skimming sample and 30 complete to mostly complete plastomes. The coalescent-based species tree, as well as the plastome tree, shows stable resolution at the backbone of the African *Sesuvium* clade. An annual C_3_ species, here newly described, is sister to a C_4_ clade that is subdivided into two subclades, one comprising the annual *S. hydaspicum* (incl. *S. nyasicum*) and the other three perennial species, *S. crithmoides*, *S. congense,* and *S. sesuvioides*. Within each of these two subclades, high gene tree discordance, mainly driven by gene tree estimation error, was found.

## Introduction

*Sesuvium* L. belongs to the ice plant family (Aizoaceae), subfamily Sesuvioideae, and comprises 18 species (Bohley & al., 2017; Sukhorukov & al., 2021). *Sesuvium* species are annuals or herbaceous perennials with opposite leaves that range from flat and slightly fleshy to terete and strongly succulent. The perennial species form cushions, and those that readily root at the nodes show a prostrate, mat-forming growth form. Depending on growing conditions, leaf color, shape, and size are phenotypically plastic. The leaves have characteristic, sometimes fringed stipules at the base of the short petioles. Single or few bracteolate, pedicellate, or sessile, 5-merous flowers are developed in leaf axils. The five perianth lobes are typically green on the dorsal side and white, rosé, or pink on the ventral side, and the androecium consists of five to many pink stamens. The 2–5-carpellate ovary ripens to a capsule that releases few to many reddish-brown to black, sculptured, or smooth seeds (Bohley & al., 2017; Sukhorukov & al., 2021). The widespread, polymorphic sea purslane (*Sesuvium portulacastrum* L.) is probably the best-known representative of the genus (Fig. 1A). It is found in coastal habitats and salty marshlands of tropical and subtropical regions worldwide and is well-known for its pronounced stress tolerance, its desalination and phytoremediation properties as well as nutritional and restoration values (Zhang & al., 2024). A second species of the genus that lately received some attention is *S. sesuvioides* (Fenzl) Verdc. (Fig. 1B) which is the only known species that performs C_4_-like photosynthesis in combination with CAM photosynthesis when stressed (Bohley & al., 2019; Siadjeu & Kadereit, 2024). Only for these two species are chromosome counts documented (https://taux.evolseq.net/CCDB_web/) and range from 2n=16–48 in *S. portulacastrum* and 2n=16 or 32 in *S. sesuvioides*.

**Fig. 1.**
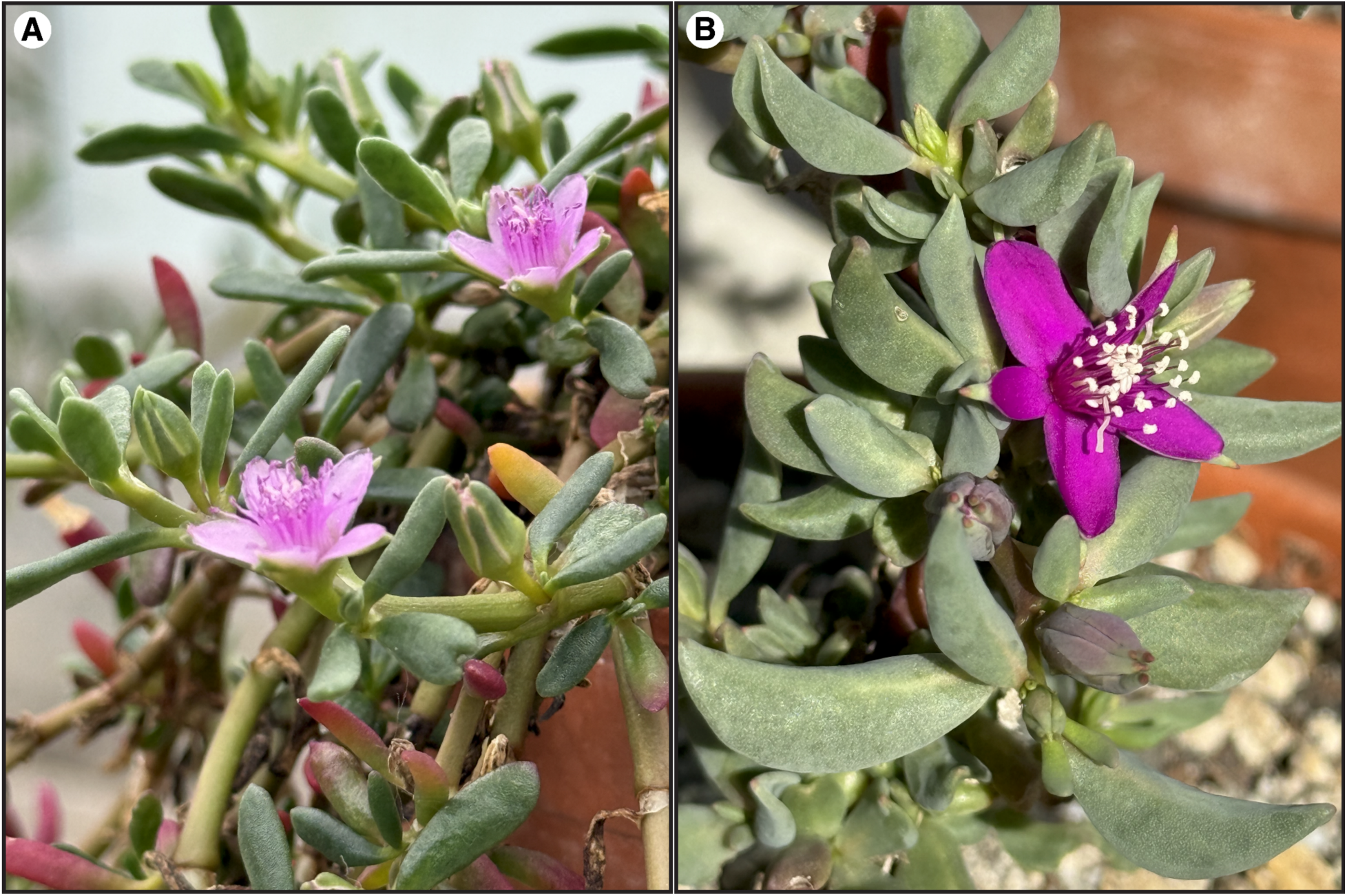
Representative species of *Sesuvium*, Aizoaceae. Flowers, buds, and leaves of the two perennial species, A. *S. portulacastrum* (L.) L. and B. *S. sesuvioides* (Fenzl) Verdc., both growing at the Munich Botanical Garden.

*Sesuvium portulacastrum* and *S. sesuvioides* represent the two main lineages of the genus consistently found in molecular systematic studies, the American (North, Central, and northern South America) lineage and the African lineage, respectively. The study of Bohley & al. (2015) was the first comprehensive molecular phylogenetic study of *Sesuvium*. It was based on five Sanger-sequenced loci and showed that the former genus *Cypselea* Turpin is nested within *Sesuvium* and likely sister to the rest of the American clade, including the widespread *S. portulacastrum*. The study of Sukhorukov & al. (2021) added several samples, especially from the Caribbean Basin, and confirmed this result.

The African lineage of *Sesuvium* has remained largely unresolved and relatively poorly sampled so far. Therefore, species boundaries, especially among the annual species, are still unclear despite recent taxonomic revisions (Bohley & al., 2017; Sukhorukov & al., 2017, 2018). Five or six species are currently accepted within the African clade, albeit species concepts do not entirely overlap between treatments (Table 1). Some diagnostic features seem clear, such as the life form and the number of bracts. In contrast, others are less helpful due to continuous intermediate phenotypes, such as the bladder cell cover on various parts of the plants and seed testa ornamentation in the case of the annual species (compare Table 1).

**Table 1.**
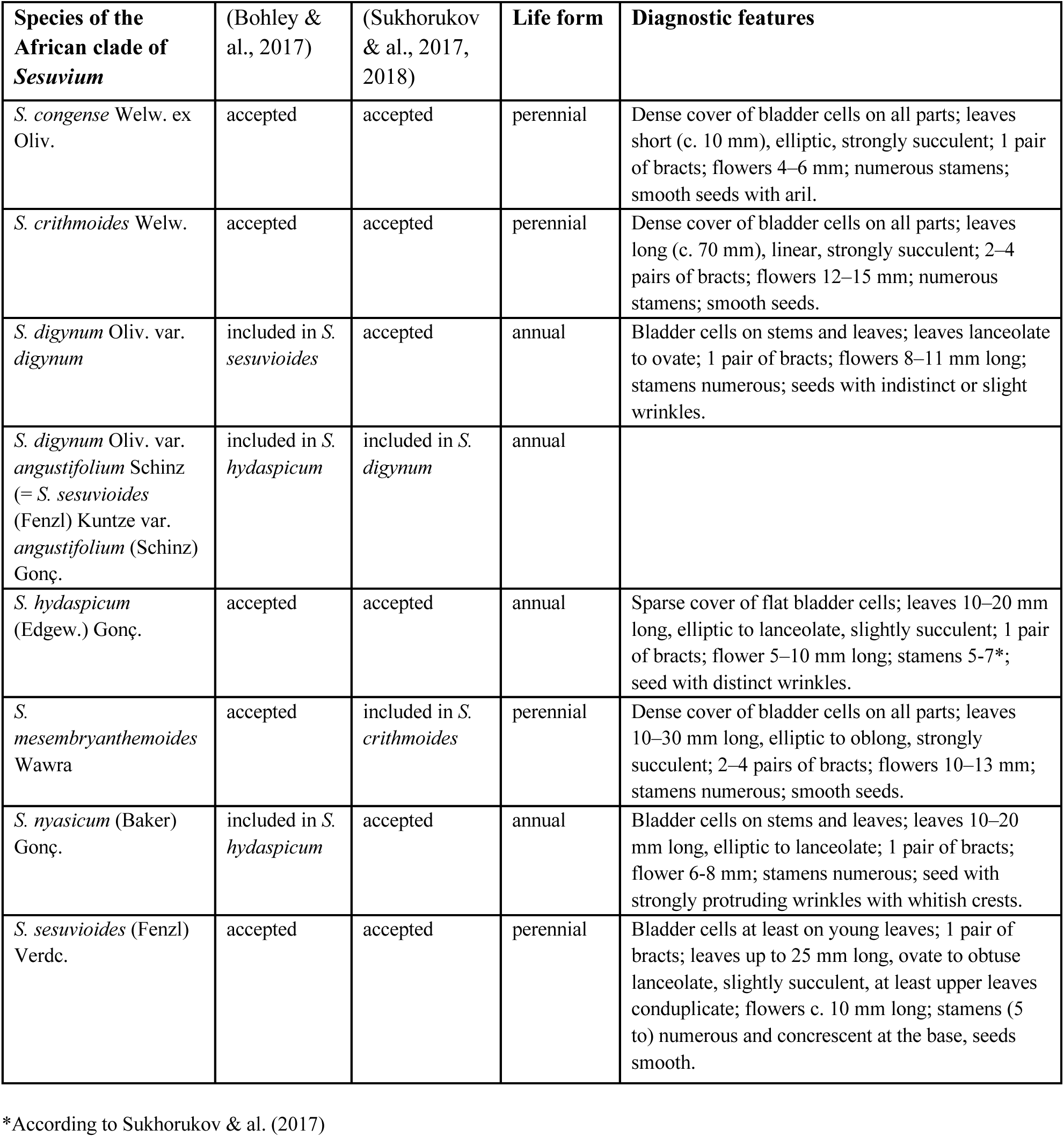
Accepted species of the African clade of *Sesuvium* in recent taxonomic treatments (Bohley & al., 2017; Sukhorukov & al., 2017, 2018) and their diagnostic features.

The Sesuvioideae are particularly interesting for studying the evolution of carbon-concentrating mechanisms (CCM). While many Aizoaceae are well-known CAM plants (Messerschmid & al., 2021; Gilman & al., 2023), most members of the subfamily Sesuvioideae are known to perform C_4_ photosynthesis (Bohley & al., 2015). Due to the structural diversity of C_4_ photosynthesis in Sesuvioideae with two biochemical subtypes (NAD-ME and NADP-ME) and several C_4_ leaf anatomical types, a multiple origin of C_4_ within the subfamily seems plausible. Within *Sesuvium,* two C_4_ origins seem to occur; one is the African lineage of *Sesuvium*, and the other is *Sesuvium humifusum* (Turpin) Bohley & G.Kadereit, a species of the American clade.

Recently, it was shown that several members of the subfamily can perform low-level CAM photosynthesis under stressful growing conditions: the C_3_ plant *Sesuvium portulacastrum* (Winter & al., 2019), the C_4_-like plant *S. sesuvioides* (Bohley & al., 2019; Siadjeu & Kadereit, 2024), and the C_4_ plant *Trianthema portulacastrum* L. (Winter & al., 2021). Only these three species have been thoroughly tested for CAM so far. An older record of possible CAM exists for *S. maritimum* (Martin & al., 1982). These occurrences of CAM in Sesuvioideae are distributed in different clades, suggesting that CAM might be overall more common in Sesuvioideae, like the rest of Aizoaceae. A possible transition from low-level CAM to C_4_ photosynthesis represents an evolutionary path to C_4_ photosynthesis that potentially differs from non-succulent C_4_ origins. Since both pathways, C_4_ and CAM, share several key enzymes, studying the integration and regulation of both pathways is particularly interesting. This was done in several studies for the genus *Portulaca* L. (Portulacaceae), which shows multiple evolution of C_4_ from ancestors with facultative CAM photosynthesis (Gilman & al., 2022 and ref. therein); however, only the model species *Portulaca oleracea* L. has been studied intensively at the transcriptomic level (Ferrari & al., 2020a, 2020b, 2021) and a more recent study by Ferrari & al. (2022) reveals candidate regulatory genes in the context of facultative C_4_–CAM photosynthesis to be validated and tested in other plants groups. The Sesuvioideae would be the ideal group to do this because of its diversity of CCM types, allowing comparative genomic studies of closely related species with diverging CCM types at a larger scale in the future.

However, since a well-resolved phylogeny is still missing, we here aim to use a deep genome skimming approach and herbarium samples to clarify the phylogenetic relationships within the African clade of *Sesuvium*. Initially, genome skimming was used as an easy way to acquire and extend the traditionally used Sanger-sequenced markers, such as ribosomal markers and chloroplast and mitochondrial markers (Straub & al., 2012). Although not always successful (e.g., Fonseca & Lohmann, 2020), it became evident that this method of shallow, random sequencing of gDNA had the potential also to deliver nuclear loci fast and at low cost, and respective pipelines for data assembly were published recently (Pouchon & al., 2018; Reginato, 2022; Pezzini & al., 2023). An improved phylogeny of the African *Sesuvium* clade is not only essential for further studies on the evolution of CCMs in this group but also serves to reassess its systematics.

## Materials and Methods

### Taxon Sampling

Taxon sampling was based on the phylogeny of Sesuvioideae from Bohley & al. (2015), and similar accessions were selected. Our focus was on the unresolved African clade and was therefore sampled in depth (19 samples representing all species of Table 1 except *S. mesembryanthemoides*, including multiple samples of *S. sesuvioides;* suppl. Table S1). Emphasis was placed on achieving a good sampling of widespread taxa. Furthermore, four species were selected from the American clade (incl. *S. portulacastrum* and the C_4_ species *S. humifusum*). A representative sampling of the remaining Sesuvioideae was also included: two species of *Zaleya* (both C_4_), five species of *Trianthema* (one C_3_, four C_4_), and two representatives of the genus *Tribulocarpus* (both C_3_). Additionally, the transcriptomes of *S. humifusum, S. portulacastrum, S. verrucosum*, *Trianthema portulacastrum, Zaleya pentandra,* and two other members of Aizoaceae (*Mesembryanthemum crystallinum* and *Delosperma echinatum*) were also included.

The genomes of *Beta vulgaris* L., *Spinacia oleracea* L. (both Amaranthaceae), and *Dianthus caryophyllus* L. (Caryophyllaceae), and the transcriptome of *Stegnosperma halimifolium* Benth. (Stegnospermataceae) were selected as outgroups from Caryophyllales. All deep genome skimming data were newly generated as described below, while genome and transcriptome assemblies were taken from Yang & al. (2015, 2018).

### DNA Extraction, Library Preparation, and Sequencing

Genomic DNA samples were available from the DNA bank (LMU, Systematics, Biodiversity and Evolution of Plants in Munich). The Qiagen DNeasy Plant Mini Kit (QIAGEN, Venlo, Netherlands) was used to isolate genomic DNA from the herbarium material of three additional samples following the manufacturer’s protocol. No RNase treatment was performed, and the final elution was done in 100µl 10mM Tris-HCl. Genomic DNA was quantified using a Qubit 4 Fluorometer with a dsDNA HS Assay Kit (Thermo Fisher Scientific, Waltham, Massachusetts, USA) and qualified by gel electrophoresis using a 0.8 % agarose gel in TAE and 30 min at 120V, 20-50 ng DNA sample. DNA samples that were not degraded required additional fragmentation to achieve an average target size of 350 bp. This was done through sonication using a Covaris M220 Focused-ultrasonicator (Covaris, Woburn, Massachusetts, USA). Library preparation was performed using the NEBNext Ultra II DNA Library Prep Kit for Illumina (New England Biolabs, Ipswich, Massachusetts, USA) according to the manufacturer’s protocol but using only half the recommended volumes to reduce the cost per sample. The protocol includes an enzymatic end-prep and adapter ligation followed by a double-sided size selection (0.26× and 0.08× volume beads). All libraries derived from highly degraded DNA were treated with the alternatively suggested clean-up using 0.9× volume beads. For multiplexing the NEBNext Multiplex Oligos for Illumina Set 2, 96 Unique Dual Index Primer Pairs (New England Biolabs, Ipswich, Massachusetts, USA) were used in a 9-cycle indexing PCR. Libraries were quantified using a Qubit Fluorometer (dsDNA HS Assay Kit) and qualified on an Agilent 4150 TapeStation System with HSD1000 ScreenTapes (Agilent Technologies, Santa Clara, California, USA). The first library preparation attempt failed in seven cases, most likely due to secondary metabolites in the DNA extracts. These libraries, with a final molarity below 10nM, were re-amplified in a subsequent PCR. All libraries were normalized to 7nM and pooled. Pre-made libraries were sequenced at Novogene UK (Novogene Company Limited, Cambridge, United Kingdom).

### Nuclear Tree-based Orthology Loci Reference Building

We inferred orthologous loci across Aizoaceae using publicly available transcriptome and genome data. All Aizoaceae transcriptome assemblies and their corresponding translations to coding sequences (CDS) and outgroup reference genomes CDS were taken from Yang & al. (2015, 2018). Seven species from Aizoaceae (including two *Sesuvium* species) and four outgroups were included (suppl. Table S2). Homology inference was done on CDS using reciprocal BLASTN searches, followed by orthology inference using the tree-based ‘monophyletic outgroup’ (MO) approach from Yang & Smith (2014) and following Morales-Briones & al. (2021). First, transcriptome CDS were reduced with CD-HIT v.4.8.1 (-c 0.99 -n 10; [Fu & al., 2012]). Then, BLASTN searches were carried out with an *E-*value cutoff of ten and max_target_seqs set to 100. BLASTN hits were filtered with a minimal fraction of 0.4. Putative homologs were then clustered with MCL v.14-137 (van Dongen, 2000) with a minimum minus log-transformed *E* value cutoff of five and an inflation value of 1.4, and only clusters with at least seven taxa were retained. Individual homolog clusters were aligned using MAFFT v.7.490 (Katoh & Standley, 2013) with settings ‘–genafpair –maxiterate 1000.’ Aligned columns were cleaned to have no more than 90% missing data using Phyx (Brown & al., 2017). Homolog trees were inferred using RAxML v.8.2.12 (Stamatakis, 2014) with a GTRCAT model and no node support. Monophyletic and paraphyletic tips from the same species were pruned from the homolog trees, keeping only the tip with the highest number of characters in the trimmed alignment. Outlier long tips were detected and removed using TreeShrink v.1.3.9 (Mai & Mirarab, 2018) with the ’per-gene’ mode and a false positive error rate threshold (α) of 0.05 while excluding the outgroups from being removed. Orthology inference was carried out keeping only ortholog groups with at least seven taxa while setting all Aizoaceae species as ingroups and *Beta vulgaris*, *Dianthus caryophyllus*, *Spinacia oleracea*, and *Stegnosperma halimifolium* as outgroups. The MO approach filters for clusters with outgroup taxa, which are monophyletic and single-copy, filtering for single- and low-copy loci. It then roots the tree by the outgroups, traverses the rooted tree from root to tip, and retains the branch with more taxa when gene duplication is detected. If no taxon duplication is detected, we set the MO approach to output a one-to-one ortholog while also checking for the outgroup taxa being monophyletic and single-copy. A total of 9,151 orthologous loci were obtained, including 6,369 one-to-one orthologs. We wrote FASTA files from all orthologs, retaining only sequences from Aizoaceae, to use as references for subsequent locus assembly from genome skimming data. The references for each locus consisted of at least three Aizoaceae sequences, including one species of *Sesuvium,* for 9,047 loci.

### Nuclear Read Processing and Assembly

We checked genome skimming raw read quality using FastQC v.0.12.1 (Andrews, 2010) and MultiQC v.1.21 (Ewels & al., 2016). Then, PCR duplicates were removed with ParDRe v.2.1.5 (González-Domínguez & Schmidt, 2016) using default parameters. We used CAPTUS v.1.0.1 (Ortiz & al., 2024) to trim sequencing adaptors and low-quality bases, assemble contigs, and extract nuclear loci based on the Aizoaceae tree-based orthologous loci data set. For the trimming and assembly steps, we used CAPTUS’ default parameters, while for loci extraction, we set the minimum percentages of identity (--nuc_min_identity) and coverage (-- nuc_min_coverage) to 75% and 50%, respectively. Along with the genome skimming data, we also extracted the Aizoaceae loci from the primary transcript of the reference genome of *Sesuvium sesuvioides* (Siadjeu and Kadereit, unpublished data). Finally, we wrote CDS FASTA files for each locus while keeping all potential paralog copies (-m NUC -f NT --max_paralogs -1 --collect_only).

### Phylogenetic Analyses

For phylogenetic analyses, we merge the corresponding sequences for each locus from the genome skimming assembled data set and from the transcriptome and genome tree-based orthologs to have a combined data set of 43 samples (39 Aizoaceae and four outgroups; suppl. Tables S1 and S2). Individual loci were aligned with the OMM_MACSE v.12.01 pipeline (Scornavacca & al., 2019). Aligned columns were cleaned to have no more than 90% missing data using Phyx. Homolog trees were inferred using IQ-TREE v.2.2.2.6 (Minh & al., 2020) using an extended model selection (Kalyaanamoorthy & al., 2017) and no clade support. Monophyletic and paraphyletic masking, outlier detection and removal, and orthology inference were done the same way as in the reference-building step, but retaining only orthologs with at least 30 samples present. Individual ortholog loci were aligned with OMM_MACSE, cleaned with Phyx, and trees were inferred with IQ-TREE using extended model selection and 1,000 Ultrafast bootstrap (UFBoot; Hoang & al., 2018) replicates for node support. Ortholog gene trees were then used to infer a coalescent-based species tree with ASTRAL-IV v.1.19.4.5 (Zhang & al., 2025) using local posterior probabilities (LPP; Sayyari & Mirarab, 2016) to assess clade support and an average gene sequence length of 1,500 for branch length estimation in substitution units with CASTLES-2 (Tabatabaee & al., 2023; integrated with ASTRAL-IV).

### Assessment of Discordance: Gene Tree Estimation Error, Incomplete Lineage Sorting, and Ancient Gene Flow

We explored gene tree discordance first by calculating the number of concordant and discordant bipartitions on each node of the ASTRAL-IV species tree using Phyparts (Smith & al., 2015). We used individual gene trees with UFBoot support of at least 70% for each node. Additionally, to distinguish conflict from poorly supported branches, we performed a Quartet Sampling (QS; Pease & al., 2018) analysis using the concatenated matrix with a partition by gene (--genetrees), the ASTRAL-IV species tree, and 5,000 replicates. QS evaluates the relative support among the three possible quartet topologies for a node (Quartet Concordance, QC), the disparity between the sampled proportions of the two discordant quartet topologies (Quartet Differential, QD), and quantifies the proportion of informative quartet replicates (Quartet Informativeness, QI)

Furthermore, to disentangle the potential causes of gene tree discordance, we used the approach from Cai & al. (2021) to estimate the relative contribution of the most common sources of gene tree discordance: incomplete lineage sorting (ILS), ancient gene flow, and gene tree estimation error. The approach of Cai & al. (2021) uses a multiple regression model (Grömping, 2006) that, in turn, uses variance decomposition to assign shares of relative importance to ILS, ancient gene flow, and gene tree estimation error in generating gene tree discordance. Each regressor’s contribution is just the R^2^ from univariate regression, and all univariate R^2^-values add up to the full model R^2^ (Grömping, 2006). We estimated levels of ILS, ancient gene flow, and gene tree estimation error for each of the internal nodes of our species tree. We estimated the levels of ILS using the population size parameter theta (ϴ) following Cloutier & al. (2019). To calculate theta for each internal branch of the species trees, we first estimated a coalescent-based tree with branch lengths in coalescent units using ASTRAL-III v.5.7.1 (Zhang & al., 2018) from the final ortholog gene trees. Then, we estimated an ultrametric tree with branch lengths in mutational units (μT, where μ is the mutation rate per generation and T is the number of generations) with PAUP v4.0a (build 168; Swofford, 2002) by constraining a maximum likelihood tree search to the ASTRAL-III species tree and using a concatenated alignment of all 7,138 orthologs, a GTRGAMMA model, and enforcing a strict molecular clock. The mutational branch lengths from the constrained tree and branch lengths in coalescent units (τ = T/4Ne) from the ASTRAL-III tree were used to estimate the population size parameter theta (ϴ = μT/τ; Degnan & Rosenberg, 2009). The larger the parameter theta, the larger the gene tree discordance product of ILS. We estimated the level of ancient gene flow using the Reticulation Index (RI) from Cai & al. (2021), which uses triplet frequencies to test for deviations from the multispecies coalescent model to detect gene flow, where gene flow between non-sister species will lead to an asymmetrical frequency distribution of the two discordant minor topologies in a rooted three-taxon phylogeny. First, observed triplet frequencies are compared to a simulated null distribution using coalescent and mutational parameters estimated from the empirical data set to estimate the RI. The null distribution is generated by simulating gene trees from 100 bootstrap species trees estimated from the empirical data set. Trees are simulated using the “sim.coal.mpest” function in Phybase 2.0 (Liu & Yu, 2010). Then, the null frequency distribution of each triplet is summarized across the bootstrap replicates. For each asymmetrical triplet, the two species exchanging alleles, based on triplet frequencies, are identified. The two introgressing branches are mapped to the species tree, and all nodes on these two branches are considered involved in introgression. Finally, for each node on the species tree, the number of mapped introgression branches is counted and normalized by the total number of triplets associated with that node. The resulting percentage is the RI. To calculate the RI, we used the scripts from Cai & al. (2021); https://github.com/lmcai/Coalescent_simulation_and_gene_flow_detection) to simulate gene trees, summarize triplet frequency distribution, and map unbalanced triplets to the species tree. The input was 1) the rooted ASTRAL-III species tree with branch length in coalescent units from the previous step, 2) 100 rooted bootstrap ASTRAL-III species trees with branch length in coalescent units inferred using multi-locus bootstrapping from input bootstrap replicates of the individual final ortholog gene trees. Gene tree bootstrap replicates were obtained with IQ-TREE using extended model selection and 100 non-parametric bootstrap (BS) replicates, and 3) the 7,138 rooted ortholog gene trees. To infer gene tree estimation error levels, we first pruned the ASTRAL-III species trees to match the sample occurrence of each of the 7,138 ortholog alignments to account for missing data. Then, we used AliSim (Ly-Trong & al., 2022) in IQ-TREE to simulate one alignment of the same length, given the ortholog alignment and the corresponding pruned ASTRAL-III species tree to estimate the best model of sequence evolution and parameters for the model for each locus. We then simulated 100 alignments of the same length as the original alignment with AliSim for each locus using the estimated model and parameters from the previous step and the complete species tree. Finally, we infer trees from each simulated alignment with IQ-TREE using the same model and parameters from the simulation step. The 713,800 trees from simulated alignments were summarized in the ASTRAL-III species trees using RAxML. The node support represents the gene tree discordance generated by gene tree estimation error. The lower the support, the higher the estimation error. To estimate levels of gene tree discordance in the empirical data, we use gene concordance factors (gCF; Minh & al., 2020) in IQ-TREE using the ASTRAL-III species tree and the 100 BS replicates from each of the 7,138 ortholog gene trees as input. Finally, we decomposed the relative importance of ILS, ancient gene flow, and gene tree estimation error in generating gene tree discordance and their confidence interval over 100 bootstrap replicates, applying the LMG method (Lindeman et al. 1980) using the functions “boot.relimp” and “booteval.relimp” from the R package *relaimpo* (Grömping, 2006).

### Analyses of hybridization and population structure based on SNPs

Filtered paired-end reads from CAPTUS were mapped to the chromosome-scale reference genome of *Sesuvium sesuvioides* using the ‘mem’ algorithm of BWA v.0.7.18 (Li & Durbin, 2009). The resulting BAM files were then sorted with SAMtools v.1.18 (Danecek & al., 2021), merged with unaligned reads (MergeBamAlignment), marked for duplicates (MarkDuplicates) with GATK v.4.6.0.0 (McKenna & al., 2010), and indexed with SAMtools. Variant calling (HaplotypeCaller) was performed per sample with GATK in GVCF mode. The resulting GVCF files were merged (CombineGVCFs) and jointly genotyped (GenotypeGVCFs) with GATK. SNPs were retained (SelectVariants) and hard filtered (VariantFiltration - QD < 2.0 || QUAL < 30.0 || SOR > 3.0 || FS > 60.0 || MQ < 40.0 || MQRankSum < -12.5 || ReadPosRankSum < -8.0) with GATK to remove SNPs with low base quality or low coverage. Finally, we used VCFtoofs v.0.1.17 (Danecek & al., 2011) to filter the SNPs further to exclude sites with no alternative allele calling for any of the samples and sites with a proportion of missing data greater than 25%, while keeping only bi-allelic SNPs and SNPs that are distanced at least 100 bp, with minimum site and genotype quality scores of 30, and masking all genotypes with a depth below 5× and above 60× (--remove-indels --max-missing 0.75 --minQ 30 --min-meanDP 5 --max-meanDP 60 - -min-alleles 2 --max-alleles 2 --thin 100 --non-ref-ac-any 1).

We explore gene flow among individuals of African *Sesuvium* using the site-pattern-based *D*-statistic (ABBA/BABA) and the *f_4_* admixture ratio (Durand & al., 2011; Patterson & al., 2012) on biallelic SNPs in Dsuite v.0.5 (Malinsky & al., 2021). In a rooted quartet [(((P1,P2),P3),O)], where the outgroup O carries the ancestral allele, A, and the derived is B, the null hypothesis assumes equal frequencies of the ABBA (P2 and P3 sharing the derived allele) and BABA (P1 and P2 sharing the derived allele) patterns due to ILS and no gene flow. A significant deviation from equal frequencies is a signal of gene flow between P3 and either P1 or P2. The *f_4_* admixture ratio is the same as the *D*-statistic up to a normalization factor. We used Dsuite’s *Dtrios* to calculate D and *f_4_* for all possible trios of samples by chromosome using the ASTRAL-IV species tree as a guide, and summarized the results with *DtriosCombine*. We plotted D and *f_4_* with *plot_d.rb* and *plot_f4ratio.rb,* respectively (https://github.com/millanek/tutorials/tree/master/analysis_of_introgression_with_snp_data/). We also used *Fbranch* to calculate the *f*-branch (Malinsky & al., 2018) to disentangle correlated *f_4_* results and assign gene flow to specific, possibly internal, branches on a phylogeny (Malinsky & al., 2021). We plotted *f*-branch results with Dsuite’s *dtools.py*.

We identified differentiation axes among individuals of African *Sesuvium* with principal components analysis (PCA) using PLINK v.1.9.0-b.7.7 (Purcell & al., 2007). PCA is a parameter-free approach with no a priori population structure assumptions. We used PLINK with a window of 50 Kb, a window step size of 5 bp, and linkage pruning with an r^2^ threshold of 0.1. We also used ADMIXTURE v.1.3.0, a model-based approach, to assess population structure in a Maximum Likelihood framework (Alexander & al., 2009). ADMIXTURE fits a model of *K* population clusters to the data, fitting individuals in a way that minimizes deviation from the Hardy-Weinberg equilibrium. We ran ADMIXTURE using the same input used for the PCA, *K* = 1–10 and performed cross-validation error estimation (cv=25) to assess the most suitable value of *K* (Alexander & Lange, 2011).

### Plastome Assembly and Tree Inference

We assembled plastomes from our 30 genome skimming libraries. First, sequencing adapters and low-quality bases were removed with Trimmomatic v.0.39 (ILLUMINACLIP: TruSeq_ADAPTER: 2:30:10 SLIDINGWINDOW: 4:5 LEADING: 5 TRAILING: 5 MINLEN:25; Bolger & al., 2014). Then, we used Fast-Plast v.1.2.9 (McKain & Wilson, 2017) with default parameters to assemble full plastomes. We used all samples of Caryophyllales (-- bowtie_index) for read mapping while expanding the Fast-Plast plastome database to add four additional Aizoaceae species from available reference plastomes, including two species of *Sesuvium* (suppl. Table S3). When the resulting plastomes did not comprise a single contig (13 out of 30), filtered contigs from SPAdes v.3.15.3 (Bankevich & al., 2012) were mapped to the reference plastome of *S. sesuvioides* (NC070250) using Geneious Prime v.2024.0 (https://www.geneious.com) to produce oriented and contiguous contigs with missing regions masked with ‘?.’ Additionally, we assemble plastomes from RNA-seq libraries of seven transcriptome samples used during the Aizoaceae tree-based orthology inference (suppl. Table S3). Raw reads were obtained from the NCBI Sequence Read Archive (SRA) (suppl. Table S3) and processed identically to the genome skimming data. For phylogenetic analyses, we included the plastome from the reference genome of *S. sesuvioides* and five GeneBank reference plastomes (*Mesembryanthemum crystallinum* and four outgroups; suppl. Table S3) to match the sampling in the nuclear data set. All plastomes, with one inverted repeat removed, were aligned with MAFFT, and columns with more than 50% missing data were trimmed with Phyx. An ML tree was inferred with IQ-TREE with automated extended model selection and 200 non-parametric bootstrap (BS) replicates. Additionally, we evaluated branch support with QS using 1,000 replicates.

### Carbon isotope value measurement

For two leaf tissue samples from the herbarium specimen “sesu130” (voucher info see suppl. Table 1), the delta C value was measured following the procedure described in (Messerschmid & al., 2021). Three technical replicates were measured for each tissue sample. Total carbon content and δ^13^C values were analyzed at the Stable Isotope Ecology Lab at the Department of Environmental Sciences, University of Basel.

## Results

### Genome Skimming, Nuclear Loci Assembly, Orthology Inference, and Plastome Assembly

The number of genome skimming raw paired-end reads ranged from 19.81 (*Sesuvium* sp. cf. sesu81) to 35.02 million (*Trianthema ceratosepala* sesu73), with an average of 25.70 million. After deduplication and cleaning, the average read number was reduced to 21.51 and 21.27 million, respectively. This would represent a ∼10× genome coverage, assuming a haploid reference genome assembly of 659.10 Mbp for *S. sesuvioides* (Siadjeu and Kadereit, unpublished data). The number of recovered loci ranged from 3,239 (*Trianthema sheilae* sesu51) to 9,138 (*S. portulacastrum* sesu74) out of 9,151, with an average of 8,505. The number of putative paralogs ranged from 191 (*T. sheilae* sesu51) to 6,000 (*S. portulacastrum* sesu74), with an average of 2,051. The number of orthologs with at least 30 species for the full data set was 7,138, with a mean of 6,182 orthologs per species. From these, 3,392 were one-to-one orthologs. The number of orthologs per genome skimming sample ranged from 2060 (*T. sheilae* sesu51) to 7,021 (*Zaleya pentandra* sesu40), with an average of 6,289. The concatenated matrix for the full data consisted of 10,461,324 aligned columns with a character occupancy of 71.55% (suppl. Table S4). We assembled 17 complete plastomes (single contig) and 13 partial (mostly complete) plastomes from genome skimming data, and only partial plastomes from transcriptome data. Complete plastome sizes ranged between 155,546 bp (*S. humifusum* sesu75) and 156,187 bp (*S. portulacastrum* sesu74). Partial plastome sizes (without IRA) for genome skimming data ranged from 123,009 (*Z. redimita* sesu95) to 130,444 (*S. sessile* sesu61) and 62,659 (*Z. pentandra* trp) to 92,783 (*S. verrucosum* trp) for the transcriptome data. The final plastome alignment (without IRA) was 129,915 bp with an occupancy of 91.10%.

### Phylogenetic Analyses and Discordance

The coalescent-based species tree showed LPP values above 0.98 throughout the tree (Fig. 2A). The plastome tree showed high support (BS > 95) along the backbone of the tree and most branches within the African clade of *Sesuvium* were notably short, where it does not show any clear species pattern (Fig. 2B). The following relationships received high support in the nuclear and plastome data sets (Fig. 2): *Sesuvium* is monophyletic and sister to *Zaleya*; within *Sesuvium*, the African and American clades are sister to each other; within the American clade, the C_4_ species *S. humifusum* (former *Cypselea humifusa*) is sister to the remaining American species (all C_3_ species); within the African clade (all C_4_ or C_4_-like species), one sample (sesu130) is sister to all remaining African *Sesuvium* samples. This sample, based on δ13C values, is a C_3_ plant, and it is morphologically distinct from the described species (see below). It represents a new annual species of *Sesuvium* (see Taxonomic Treatment).

**Fig. 2.**
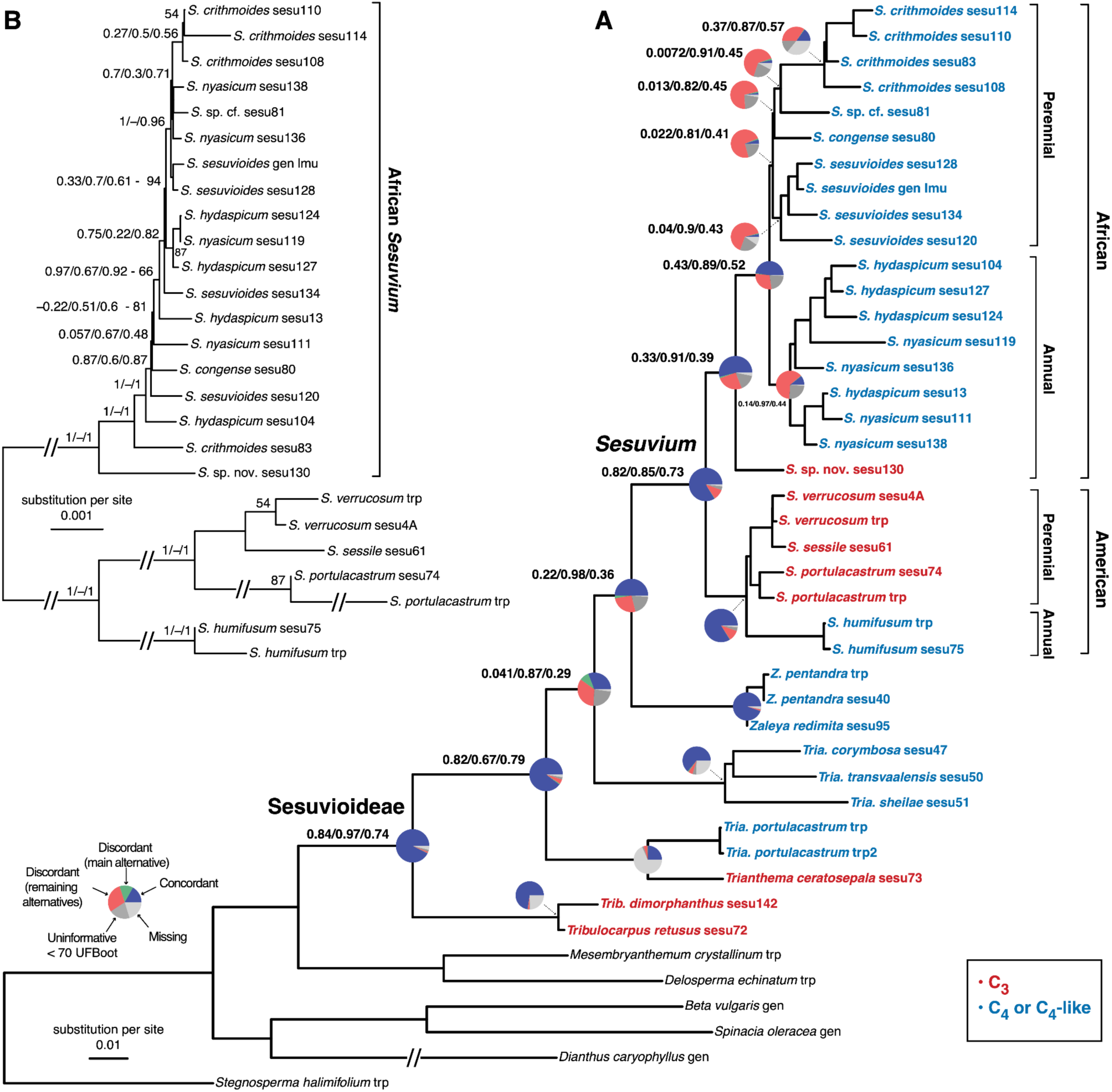
A. Coalescent-based species tree of Sesuvioideae inferred with ASTRAL-IV from 7,138 orthologous gene trees. Tip labels of Sesuvioideae are colored by photosynthetic pathway, C_3_ in red and C_4_ or C_4_-like in blue. All nodes have a local posterior probability (LLP) of at least 0.98. Pie charts for main nodes represent the proportion of gene trees that support that clade (blue), the main alternative bifurcation (green), the remaining alternatives (red), the remaining alternatives with (conflict or support) with Ultrafast bootstrap (UFBoot) lower than 70% (dark gray), and missing data (light gray). Quartet Sampling (QS) scores are shown next to the pie charts. QS scores: Quartet concordance/Quartet differential/Quartet informativeness. Branch lengths in number of substitutions per site (scale bar). *Dianthus caryophyllus* branch is shortened with a broken segment (//) for illustration purposes (see suppl. Figs. S1–S3 for original branch lengths, UFBoot, pie charts, and QS scores for all nodes). B. Maximum likelihood phylogeny of Sesuvioideae inferred with IQ-TREE from whole plastomes depicting only *Sesuvium.* QS scores next to backbone nodes. All nodes have a non-parametric bootstrap (BS) of at least 95 % unless noted next to QS scores or nodes. Long branches were shortened with a broken segment (//) for illustration purposes (see suppl. Figs. S4–S5 for original branch lengths, full phylogeny of Sesuvioideae, and UFBoot and QS scores for all nodes. Branch lengths in number of substitutions per site (scale bar).

Among the remaining African samples, the resolution of the plastome tree is low, and the gene tree discordance of the nuclear and plastome trees is high (Fig. 2A-B). The coalescent-based species (Fig. 2A) resolves two sister clades of annual versus perennial species, respectively. The ‘perennial’ clade includes *S. crithmoides*, *S. congense,* and *S. sesuvioides*. The ‘annual’ clade includes samples of *S. hydaspicum* and *S. nyasicum*. These two clades are supported by a low fraction (332 out of 5318 for the ‘perennial’ and 755 out of 4526 for the ‘annual’ clade) of concordant gene trees (Fig. 2 and suppl. Fig. S2). The QC score within both clades lies for most nodes between 0 and 0.25, and the QD score is close to 1, which means the two discordant alternatives are equal in frequencies (Fig. 2 and suppl. Fig. S3). This shows that the two clades are stable despite the gene tree discordance.

Gene tree discordance estimated using gCF was particularly high for all nodes of African *Sesuvium* (Fig. 3 and suppl. Fig. S6). The multiple regression approach showed that the largest contributor to gene tree discordance is gene tree estimation error (84.5%) and not ILS (11.7%) or ancient gene flow (3.7%; Fig. 3). While the regression was done on the entire data set, African *Sesuvium* had the nodes with the most gene tree estimation error, ILS, and in part, ancient gene flow (suppl. Fig. S6) and despite the nodes around *Trianthema* showing similar patterns to African *Sesuvium,* the latter is the one driving most of the signal (suppl. Fig. S6). Interestingly, gene tree estimation error negatively correlated (*R* = − 0.17; *p* = 0.3) with the estimate of ancient gene flow, while it was positively correlated with ILS (*R* = 0.39; *p* = 1.3×10^-2^) and gene tree discordance (*R* = 0.54; *p* = 3.2×10^-4^). ILS and ancient gene flow had a positive correlation (*R* = 0.45; *p* = 3.4×10^-3^), suggesting that RI might not have effectively distinguished these two processes, but still, their contribution is significantly smaller than the one from gene tree estimation error (suppl. Fig. S7).

**Fig. 3.**
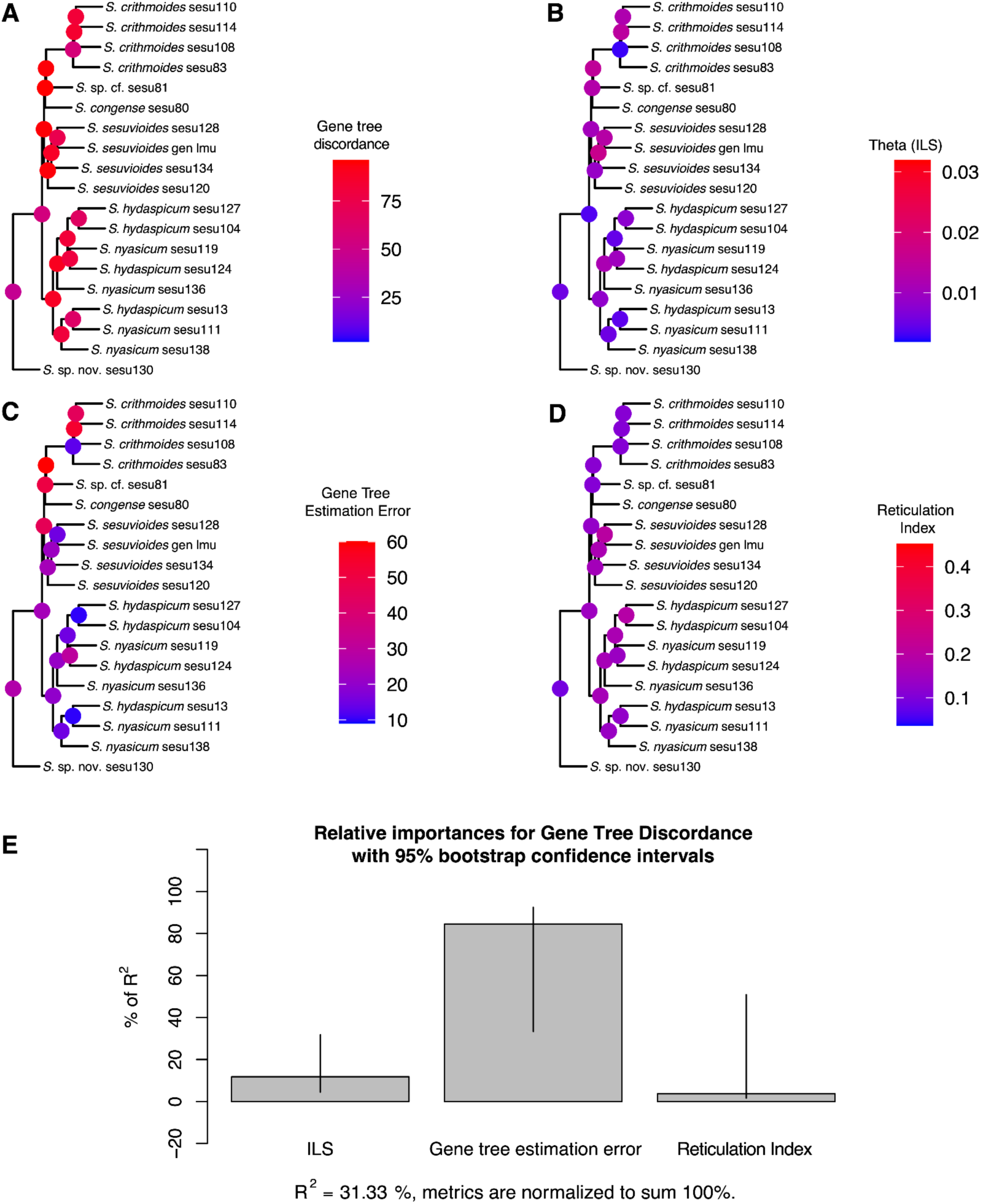
Relative contribution of the most common sources of gene tree discordance (A): Incomplete lineage sorting (ILS; B), gene tree estimation error (C), and ancient gene flow (D). A. Gene tree discordance is shown as inverse concordance factors (gCF). The higher the value, the higher the discordance. B. ILS is depicted by the population size parameter theta. Higher theta values represent higher ILS. C. Gene tree estimation error shown as the inverse node support from simulated trees. The higher the value, the higher the estimation error. D. Ancient gene flow depicted by the Reticulation Index (RI). The higher the RI, the higher the gene flow. A–D showing only African *Sesuvium.* Values for all nodes in the complete data set are shown in suppl. Fig. S6. E. Relative importance of ILS, gene tree estimation error, and ancient gene flow in generating gene tree discordance. The percentages are estimated based on the LMG method. The 95% confidence intervals are represented by bars. The contribution of the regressor was estimated using all nodes from the complete tree (suppl. Fig. S6).

### SNP analyses of population structure and hybridization

The PCA analysis of SNP data supported the overall topology described above. Most of the differentiation is shown by PC1 and PC2 (suppl. Fig. S8) and revealed strong separation of the new *Sesuvium* species (sample sesu130) from the remaining African *Sesuvium* (Fig. 4). Furthermore, the ‘perennial’ and ‘annual’ clades formed distinct clusters. While the cluster of samples from perennial species showed a substructure that separated the species in that clade (*S. sesuvioides*, *S. congense,* and *S. crithmoides*), this was not the case for the samples of the two annual species, *S. hydaspicum* and *S. nyasicum*, which showed a clear overlap between the two species (Fig. 4). The cross-validation error estimation showed that *K = 6* is the optimal number of populations (suppl. Fig. S9). The ADMIXTURE analyses are congruent with the PCA. Sample sesu130 differs from the rest of African *Sesuvium*. *Sesuvium sesuvioides*, *S. congense,* and *S. crithmoides* are independent populations, while *S. hydaspicum* and *S. nyasicum* showed clear overlap. Sample sesu81, which was identified as *S. sesuvioides* in Bohley & al. (2017), clustered with other representatives of that species in the PCA and ADMIXTURE analyses (Fig. 4), while it was placed as sister to *S. crithmoides* in the coalescent-based species tree (Fig. 2).

**Fig. 4.**
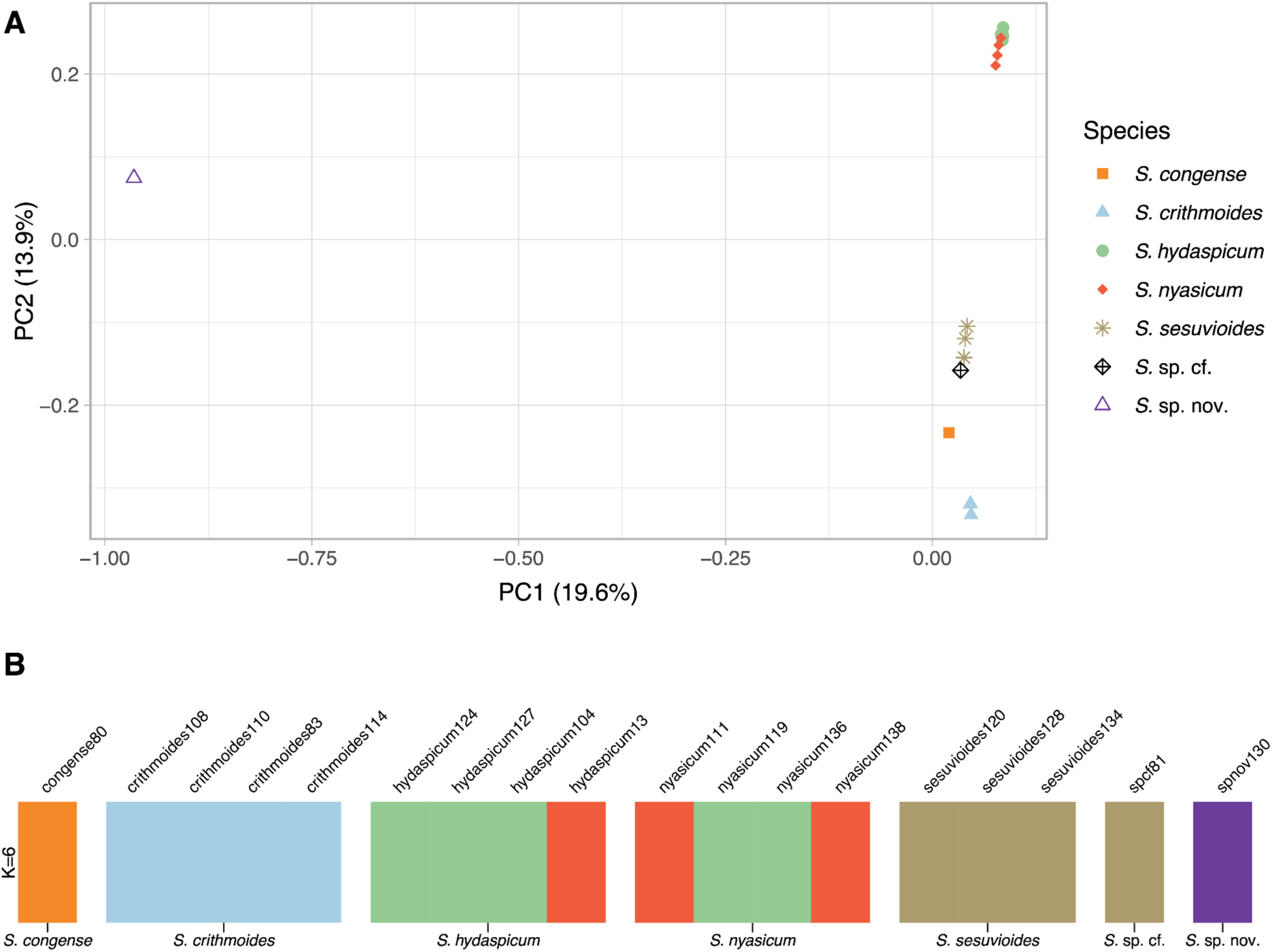
A. Principal component analysis (PCA) of African *Sesuvium* based on 330,414 genome-wide SNPs inferred with PLINK, showing the two main axes of differentiation. B. ADMIXTURE model-based clustering analysis of African *Sesuvium* based on 330,414 genome-wide SNPs, showing the optimal *K*=6 (see suppl. Fig. S10 for clustering with *K* = 2–10).

The *D*-statistic, *f_4_* admixture ratio, and *f*-branch analysis on biallelic SNPs indicate limited gene flow between species of African *Sesuvium* (Fig. 5). The *D*-statistic (Fig. 5A) showed gene flow between all samples of *Sesuvium sesuvioides* with *S. hydaspicum* and *S. nyasicum.* Similarly, the unassigned sample, sesu81, showed signals of gene flow with *S. hydaspicum, S. nyasicum,* and *S. congense.* Lastly, a signal of gene flow was detected between *S. congense* and an individual of *S. sesuvioides* (sesu120). These patterns are significantly less pronounced in the *f_4_* admixture ratio (Fig. 5B). The *f*-branch showed a strong signal of hybridization only between one sample of *S. hydaspicum* (sesu124) and *S. nyasicum* (sesu129) and might explain the placement of the latter as sister of most *S. hydaspicum* samples in the coalescent-based species trees (Fig. 2A) and their sister relationship on the plastome tree (Fig. 2B). Sample sesu81 showed signals of hybridization with *S. hydaspicum, S. nyasicum,* and *S. sesuvioides,* while *S. sesuvioides* (sesu120) with *S. hydaspicum, S. nyasicum,* which is in part congruent with the *D*-statistic and *f_4_* admixture ratio, but the signals are weaker in all cases.

**Fig. 5.**
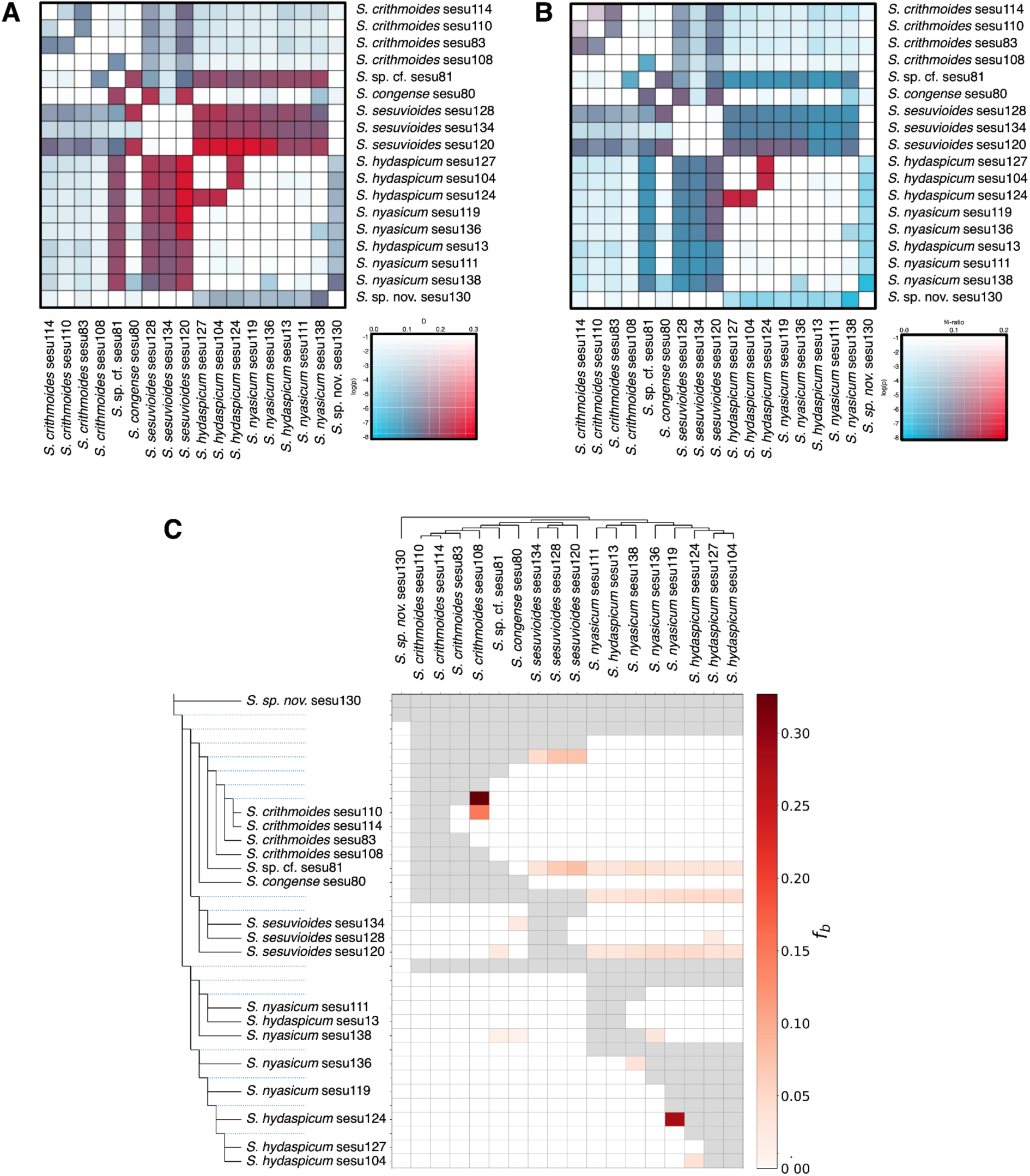
*D*-statistic (A) and *f_4_* admixture ratio (B) among members of African *Sesuvium* calculated from ABBA/BABA site patterns using 597,387 genome-wide SNPs. Colors correspond to statistical support (light to dark) and magnitude (blue to red) of the statistics. C. *f*-branch statistics for members of African *Sesuvium.* The species tree used is shown along the x- and y-axes. Dotted blue lines in the tree on the y-axis represent ancestral lineages. Darker colors depict increasing evidence of gene flow between lineages. Grey squares represent trio combinations incompatible with the species tree.

### Carbon isotope

The δ^13^C value of plant tissues allows for the distinction of different photosynthetic pathways (Cernusak & al., 2013). For a C_3_ plant where Rubisco is the only carboxylating enzyme within the photosynthetic cycle, the δ^13^C value usually lies around -27.1‰ (O’Leary, 1981), but might vary considerably from -20 to -37.5‰ due to various factors influencing isotope fractionation (Farquhar & al., 1989; Winter & Holtum, 2002). δ^13^C values less negative than -20‰ indicate the presence of a carbon concentrating mechanism in which a large proportion of carbon is fixed by phosphoenolpyruvate carboxylase (see Messerschmid & al., 2021). For specimen “sesu130”, we measured δ^13^C values of -25.8‰ and -25.5‰, which indicate predominant C_3_ photosynthesis and exclude C_4_ photosynthesis and constitutive CAM photosynthesis. For the remaining species of *Sesuvium,* we adopted the δ^13^C values measured or compiled in previous studies (Bohley & al., 2015).

## Discussion

Next-generation sequencing technologies and new bioinformatic tools have greatly advanced molecular systematics by refining and improving previous phylogenetic trees based on a few Sanger-sequenced markers (Guo & al., 2023). Transcriptome sequencing (RNA-seq), target enrichment, restriction-site associated DNA sequencing (RAD-seq), and genome skimming are the most widely used applications nowadays. A recent study by Mo & al. (2025) on the phylogenomics of *Rhododendron* shows the great potential of deep genome skimming, also when historical material is used and shallow taxonomic levels are investigated. Both points apply to our study of *Sesuvium*, and we found that, also for this group, deep genome skimming provides large amounts of informative data and proves a highly efficient and economic approach.

Previous molecular phylogenetic analyses based on a low number of Sanger-sequenced markers resolved only the deeper nodes within *Sesuvium* with the sister group relationship of an African and American clade and revealed the nested position of the former genus *Cypselea* in the American clade (Bohley & al., 2015; Sukhorukov & al., 2021). This led to the inclusion of *Cypselea* into *Sesuvium* (Bohley & al., 2017). These results and taxonomic conclusions are confirmed here based on data retrieved from 7,138 orthologous genes sequenced with a deep genome skimming approach. The data sets of the earlier studies were not sufficient to reveal internal phylogenetic relationships within the African or the American clade. Our approach generates sufficient phylogenomic signal to further resolve the relationships among the densely sampled species of the African clade. However, we found large amounts of gene tree discordance that seems to be driven mainly by gene tree estimation error and, to a much lesser degree, by ILS and hybridization.

We discovered a new lineage within the African *Sesuvium* clade, which comprises a new C_3_ species (*S. ligulifolium*, spec. nov.) and is sister to the remaining African species, which are either C_4_-like (*S. sesuvioides*) or C_4_ species (Sankhla & al., 1975; Bohley & al., 2015), and references therein). This finding supports the parallel origin of C_4_ photosynthesis of the same biochemical subtype (NADP-ME) in the African and American clade and makes a C_4_ ancestor for *Sesuvium* and a reversal to C_3_ within the American clade unlikely, especially as the sister genus *Zaleya* shows a different biochemical C_4_ subtype (NAD; compare Bohley & al., 2015, 2019). This underlines once more the importance of dissecting complex traits such as C_4_ photosynthesis into several traits for reliable character state reconstructions (Kadereit & al., 2014; Bohley & al., 2015). Furthermore, there is no indication in the molecular data of reticulation or hybridization events being involved in the evolution of C_4_ photosynthesis within the African *Sesuvium* clade, as has been shown, for example, in *Flaveria* (Morales-Briones & Kadereit, 2023). In contrast to *Flaveria* (see, for example, Adachi & al., 2023), there are no species with proto-Kranz or C_2_ photosynthesis phenotypes, but only the C_4_-like phenotype of *Sesuvium sesuvioides* (Bohley & al., 2019). Therefore, a more detailed investigation of the C_4_ evolution of this presumably relatively young C_4_ lineage might reveal an alternative evolutionary route to C_4_ than in *Flaveria*, the current model of C_4_ evolution.

Since all species of the African clade show at least some degree of succulence, we hypothesize that not only is the C_4_-like species *S. sesuvioides* capable of low-level CAM photosynthesis (Siadjeu & Kadereit, 2024), but also the others. If this assumption holds, the African clade of *Sesuvium* also represents a particularly interesting group for studying the evolutionary integration and concerted change of low-level CAM, C_3,_ and C_4_ photosynthesis. Within this clade, so far, two C_4_ leaf anatomical types, pilosoid (*S. crithmoides*) and portulacelloid (*S. congense* and *S. sesuvioides*) have been described (Bohley & al., 2015). These differ in the arrangement of tissues. The anatomical section of the terete leaves of *S. crithmoides* shows a central water storage tissue with a few vascular bundles in the centre and a peripheral chlorenchyma consisting of mesophyll and Kranz cells around small peripheral vascular bundles. The leaves of *S. sesuvioides* and *S. congense* are dorsi-ventrally organized with a chlorenchyma consisting of mesophyll and Kranz cells around small peripheral vascular bundles on the upper side and a water storage tissue (of varying thickness) without vascular bundles on the lower side. For Sesuvioideae, two more C_4_ leaf anatomical types have been described, atriplicoid and salsoloid (Bohley & al., 2015), which offer the opportunity to study CAM-C_4_ integration in the context of different anatomical features. A similar variation of C_4_ leaf types and strikingly similar leaf anatomies have been found for the other lineage that integrates low-level CAM and C_4_, the Portulacaceae (Ocampo, 2015); however, in this group, the different C_4_ leaf anatomies seem consistent in older clades.

### The Annual Species clade

Eight samples of annual *Sesuvium* were included. These represented *S. digynum*, *S. hydaspicum,* and *S. nyasicum* according to the key and descriptions provided in Sukhorukov & al. (2017). The overall resolution of this clade was low, and none of the three species could be resolved as monophyletic. The principal component analysis (PCA) based on 330,414 genome-wide SNPs revealed the lack of genetic differentiation among the samples of the annual species. The main diagnostic feature used to separate the species is the seed ornamentation of indistinct to distinct protruding wrinkles (Table 1); however, this character seems rather continuous than discrete. Taken together, these points to an aggregate of morphologically and genetically not distinct species, and therefore we suggest treating them under the name *Sesuvium hydaspicum* (Edgew.) Gonç.

### The Perennial Species clade

Ten samples of perennial *Sesuvium* were included. They represented *S. congense*, *S. crithmoides,* and *S. sesuvioides*. In contrast to the samples of annual species, those of the perennial species are well distinct from each other in the PCA (Fig. 4A), and except one sample (Sesu 81) also in the ASTRAL-IV tree (Fig. 2). Probably the most distinct among the perennial species is *S. crithmoides* with its 2 to 4 pairs of bracts (while all other species just have one pair of bracts) and by its long, strongly succulent terete leaves. *Sesuvium congense* seems genetically distinct from the other species (Fig. 4A, B); however, the inclusion of more samples is needed to test this rigorously. *Sesuvium congense* and *S. crithmoides* share a dense cover of bladder cells on all parts and highly succulent leaves, but the leaves of *S. congense* are much shorter (10 versus 70 mm in *S. crithmoides*) and elliptic. Furthermore, the flowers are also distinctly smaller (compare Table 1).

The probably most polymorphic and certainly the most widespread species of the perennial *Sesuvium* is *S. sesuvioides* (Sukhorukov et al., 2017, 2018), and likely more samples would be needed to capture the full diversity of this species. However, the five samples included form a cluster in the PCA (Fig. 4A), and four are united by several concordant gene trees (Fig. 2), indicating that this species is genetically distinct.

### Taxonomic Treatment

The five accepted species within the African clade and their synonyms

1. ***Sesuvium congense*** Welw. ex Oliv., Fl. Trop. Afr. 2, 586. 1871.

Synonyms: None.

Note: Among the three perennial species, *S. congense* is distinct by its short (c. 10 mm long) leaves. The leaves of *S. crithmoides* and *S. sesuvioides* are c.70 mm and >20 mm long, respectively.

Distribution and Habitat: Angola, from Benguela to Namibe province; coastal or inland sandy or gravelly soils.

2. ***Sesuvium crithmoides*** Welw., Apont. 586. 1858.

Synonyms: Heterotypic - *Sesuvium mesembryanthemoides* Wawra in Wawra & Peyr., Sitzungsber. Acad. Wien, Math.-Nat. 38: 564 (1860); *Sesuvium crystallinum* Welw. in Oliver, Fl. Trop. Afr. 2: 586 (1871).

Note: We follow (Sukhorukov & al., 2018) and include *S. mesembryanthemoides* and *S. crystallinum* in *S. crithmoides.* (Bohley & al., 2017) also stated that these three species are morphologically very close but did not synonymize *S. mesembryanthemoides.* Based on a review of more specimens and field observations, (Sukhorukov & al., 2018) convincingly combined all three under *S. crithmoides*. The characteristic features of this species are the 4–8 bracts and long, terete leaves, which resemble the leaves of *Crithmum maritimum* - hence the species epithet “*crithmoides*”.

Distribution and Habitat: Angola, Democratic Republic of Congo, Republic of Congo, introduced to USA (Georgia, Glynn County); subtropical, arid coasts and coastal dunes.

3. ***Sesuvium hydaspicum*** (Edgew.) Gonç. in Garcia de Orta 13: 381 (1965).

Synonyms: Homotypic - *Trianthema hydaspicum* Edgew. in J. Proc. Linn. Soc., Bot. 6: 203 (1862); heterotypic - *Sesuvium nyasicum* (Baker) Gonç. in Garcia de Orta 13: 381 (1965), *Trianthema nyasica* Baker, Bull. Misc. Inform. Kew 1897 (128–129): 268. 1897; *S. digynum* Welw. in Oliver, Fl. Trop. Afr. 2: 586 (1871); *S. digynum* var. *angustifolium* Schinz, Bull.

Herb. Boiss. 5 (Appendix 3): 74 (1897), *S. sesuvioides* (Fenzl) Kuntze var. *angustifolium* (Schinz) Gonç., *Halimus sesuvioides* (Fenzl) Kuntze var. *angustifolium* (Schinz) Hiern, Cat. Afr. Pl. 1(2): 414 (1898); *Halimus sesuvioides* (Fenzl) Kuntze var. *welwitschii* Hiern, Cat. Afr. Pl. 1(2): 414 (1898); *Trianthema polysperma* Hochst. ex Oliv., Fl. Trop. Afr. 2: 588. 1871; *Sesuvium hoepfnerianum* Schinz in Bull. Herb. Boissier 5, app. 3: 75. 1897; *Trianthema salarium* Bremekamp, Ann. Transvaal Mus.: 239. 1933.

Note: We follow (Bohley & al., 2017) and include *S. nyasicum* in *S. hydaspicum*. The main character used to differentiate the two species is the height of the wrinkles on the seed, and this feature seems to vary continuously. Samples tentatively assigned to the two former species in our analysis do not form monophyletic groups or distinct clusters based on their SNPs. According to our observations, the annual *S. digynum* (sensu Sukhorukov & al., 2017) also belongs to this species, which also has rugose to wrinkled seeds according to Sukhorukov & al. (2017).

Distribution and Habitat: In its broad circumscription, this species is widespread in arid and subtropical regions of Africa and the Arabian Peninsula, also reaching comparable habitats of Pakistan to India (type locality); however, samples from those areas were not included in any molecular analyses. Sandy or gravelly soils, disturbed areas, roadsides, and agricultural fields.

4. ***Sesuvium sesuvioides*** (Fenzl) Verdc., Kew Bull. 12(2): 349 (1957).

Synonyms: Homotypic - *Diplochonium sesuvioides* Fenzl, Nov. Stirp. Dec. 1: 58. 1839, *Halimus sesuvioides* (Fenzl) Kuntze, Revis. Gen. Pl. 1: 263. 1891; heterotypic - *Halimum sesuvioides* (Fenzl) Hiern var. *reduplicatum* Welw. ex Hiern, Cat. Afric. Pl. (Hiern) 1(2): 414. 1898, *Halimum sesuvioides* (Fenzl) Hiern var. *welwitschii* Hiern, Cat. Afric. Pl. (Hiern) 1(2): 414. 1898, *Sesuvium hoepfnerianum* Schinz var. *brevifolium* Schinz, Bull. Herb. Boiss. 5 App. III: 75. 1897.

Note: A polymorphic perennial species with a sparse to dense cover of bladder cells; in younger parts, there is often a denser cover. Typical are the flat but succulent, distinctly folded leaves. According to Bohley & al. (2017), the number of stamens varies between 5 and many. The seeds are smooth and not ornamented.

Distribution and Habitat: Angola, Cape Verde, Namibia, and the northwestern parts of South Africa (Northern Cape Province; according to Bohley et al. 2017); flats, depressions, dry riverbeds, along roads on sandy, gravely, or saline soil.

**5. *Sesuvium ligulifolium* G.Kadereit & D.F.Morales-B., *spec. nov*.**

Type: W. Giess & B. Loutit 14178, 8.3.1976, Southwest Africa, Namibia, 1916 BA Gobaub, Etosha N.P., nördlich der Wasserstelle Nuanses. Holotype M-0292626 (M!), Isotype PRE-0462875-0 (PRE). (suppl. Fig. S11).

Description - Succulent, decumbent annual herbs with dichotomous branching, 10–20 cm high, the main axis terminating with a single flower, stems show a sparse cover of bladder cells; opposite leaves equal in size, petiole c. 5mm with translucent lateral flaps, margin of the flaps entire, leaf blade c. 1.5–2 cm long and 3 mm wide, ligulate, leaves with a sparse cover of bladder cells; 1 pair of tiny, acute bracts; flowers solitary and sessile, 6–8 mm long and turbinate in bud, 5 tepals, lower half connate, upper half free and slightly hood-shaped, with a tiny but distinct globular thickening at the tip and dorsally covered in bladder cells, alternating with the free part of the tepals triangular protrusions which are more distinct in fruiting stage; ten stamens, anthers tiny, 0.5 mm long; ovary with two thread-like styles; c. 10–15 seeds, seed testa wrinkled, seed c. 1.2 mm across.

Note: The holotype contains three flowering (upper branches) and fruiting (lower branches) individuals, including roots (suppl. Fig. S11). The collectors noted that the plant shows a prostrate-ascending growth, the leaves are described as not folded (as would be typical of *S. sesuvioides*) and are 2 cm long and 3 mm wide. The open flowers are described as red-purple and estimated to be 2.5 mm across. Furthermore, the collectors describe the plants as “saftiges Kraut,” which means that the fresh leaves are distinctly succulent.

This species shares the annual life form and the wrinkled seed testa with *S. hydaspicum* (incl. *S. nyasicum*) but differs in the ligulate leaf shape and the number of stamens. Furthermore, *S. ligulifolium* is a C_3_ species while *S. hydaspicum* is a C_4_ species.

Distribution and Habitat: The type locality is currently the only confirmed collection of this species. It is on the South side of the Etosha pan, north of the Nuamses waterhole (wrongly spelled on the specimen label as “Nuanses”), approximately at -19,125 S and 16.625 E. The habitat is described by the collectors as partly flooded ground. We assume that the species might be more widespread in the area.

## Conclusion

The African *Sesuvium* clade comprises five distinct species in three subclades. The new annual species *S. ligulifolium* is sister to the remaining four species, which subdivide into a subclade of annual taxa (species aggregate S. *hydaspicum*) and a subclade of perennial species, *S. congense*, *S. crithmoides*, and *S. sesuvioides*. The molecular data point to young, evolutionarily not fully sorted lineages, which makes the whole clade interesting for studying the evolution of C_4_ photosynthesis, also in the context of the co-occurrence of weak CAM photosynthesis.

## Data Availability

Genome skimming data generated for this study can be found in the NCBI BioProject XXXXXX (see suppl. Table S1 for SRA accession numbers). Sequences of complete and partial plastomes are deposited in GenBank (see suppl. Table S1 for accession numbers). Analysis files are available from the Dryad repository https://doi.org/10.5061/dryad.XXXX

## Author Contributions

Conceptualization: GK; Formal analysis: DG, DFMB; Funding acquisition: GK; Investigation: GK, AH; Resources: GK; Supervision: GK, DFMB; Visualization: DFMB; Writing – original draft: GK, AH, DFMB; Writing – review & editing: all authors.

## Acknowledgments

We thank Dan Nelson (Univ. Basel) for measuring the carbon isotope ratio for the sample of *S. ligulifolium* and C. Siadjeu (Univ. Mainz) for providing unpublished genome data of *S. sesuvioides*. We are grateful to the staff of the Botanical State Collection Munich for help with sampling herbarium specimens and to M. Silber (LMU Munich) for extracting DNA from the partly challenging samples. Financial support came from the DFG project KA1816/7-3 to G. Kadereit and from the LMU Munich.

