## Supplemental_Figs_and_Tables for "Phylogenomics and Systematics of African *Sesuvium* (Aizoaceae)"

Fig. S1.

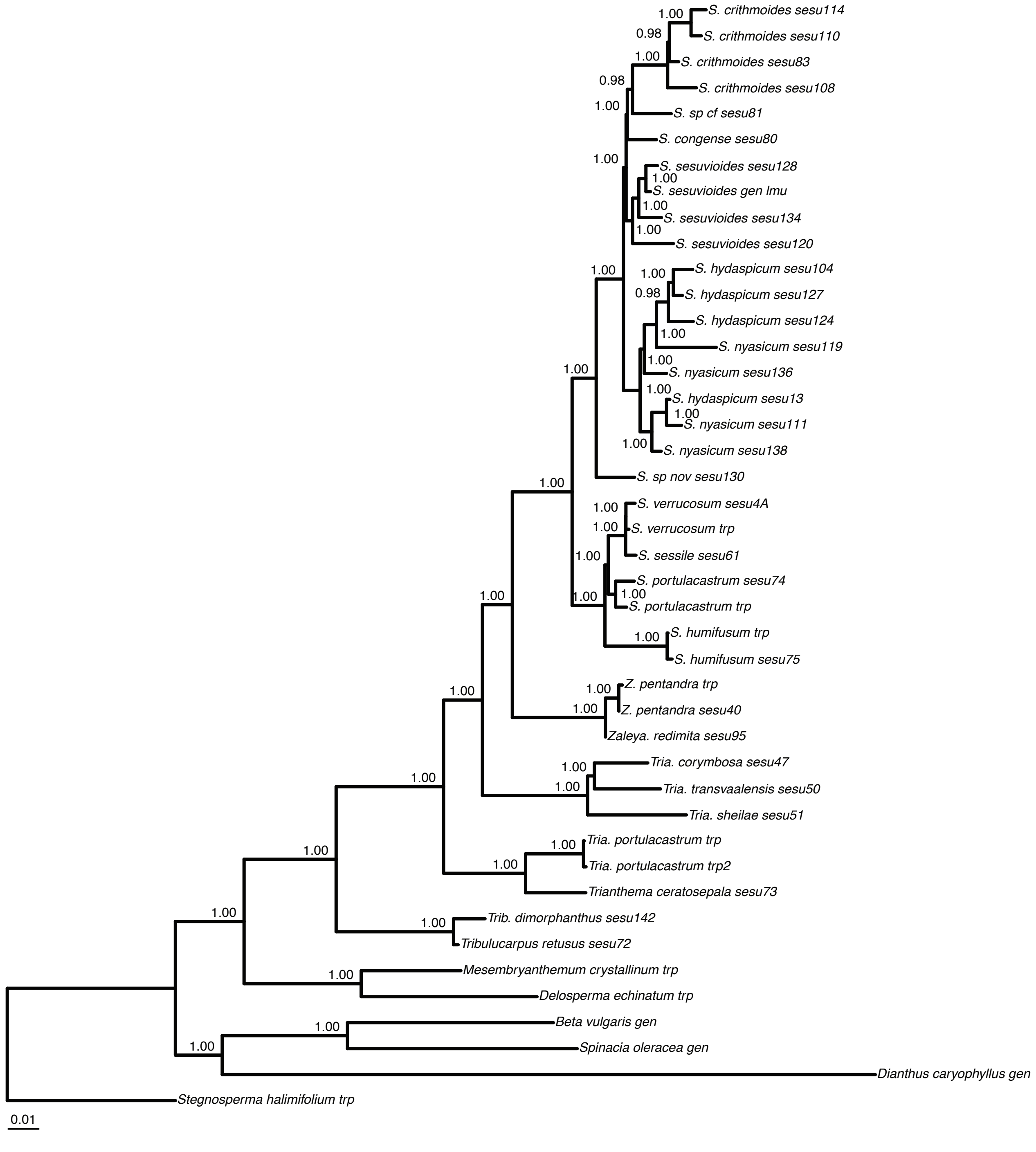
S. crithmoides sesu114
S. crithmoides sesu110
S. crithmoides sesu83
S. crithmoides sesu108
S. sp cf sesu81
S. congense sesu80
S. sesuvioides sesu128
S. sesuvioides gen lmu
S. sesuvioides sesu134
S. sesuvioides sesu120
S. hydaspicum sesu104
S. hydaspicum sesu127
S. hydaspicum sesu124
S. nyasicum sesu119
S. nyasicum sesu136
S. hydaspicum sesu13
S. nyasicum sesu111
S. nyasicum sesu138
S. sp nov sesu130
S. verrucosum sesu4A
S. verrucosum trp
S. sessile sesu61
S. portulacastrum sesu74
S. portulacastrum trp
S. humifusum trp
S. humifusum sesu75
Z. pentandra trp
Z. pentandra sesu40
Zaleya. redimita sesu95
Tria. corymbosa sesu47
Tria. transvaalensis sesu50
Tria. sheilae sesu51
Tria. portulacastrum trp
Tria. portulacastrum trp2
Trianthema ceratosepala sesu73
Trib. dimorphanthus sesu142
Tribulucarpus retusus sesu72
Mesembryanthemum crystallinum trp
Delosperma echinatum trp
Beta vulgaris gen
Spinacia oleracea gen
Dianthus caryophyllus gen
Stegnosperma halimifolium trp
0.01

Fig. S2.

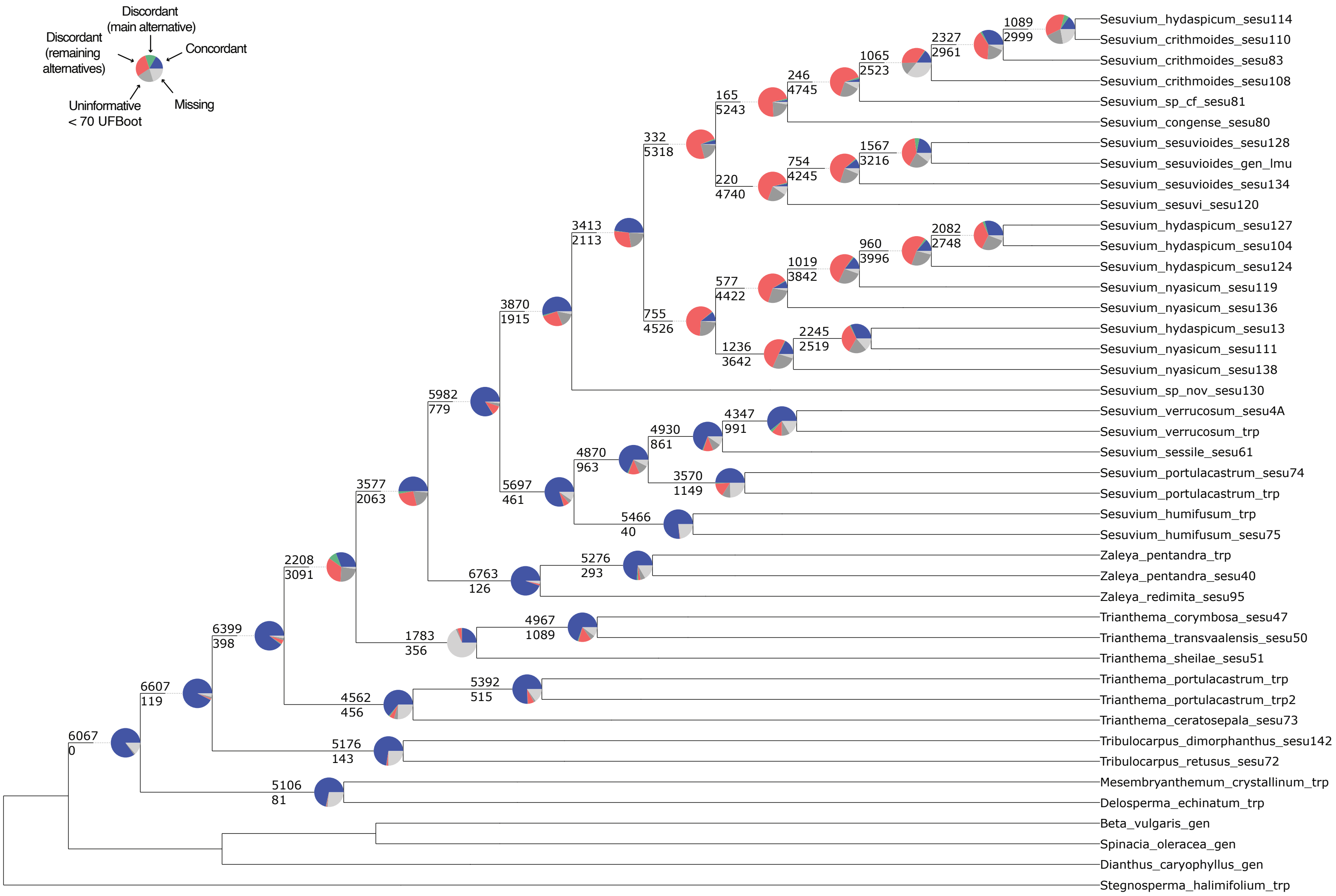

Discordant (main alternative)
Discordant (remaining alternatives)
Concordant
Uninformative < 70 UFBoot
Missing
1089
2999
2327
2961
1065
2523
246
4745
165
5243
332
5318
1567
3216
754
4245
220
4740
3413
2113
2082
2748
960
3996
1019
3842
577
4422
755
4526
2245
2519
1236
3642
3870
1915
5982
779
4347
991
4930
861
4870
963
3570
1149
5697
461
5466
40
3577
2063
5276
293
6763
126
2208
3091
4967
1089
1783
356
6399
398
5392
515
4562
456
6607
119
5176
143
6067
0
5106
81
Sesuvium_hydaspicum_sesu114
Sesuvium_crithmoides_sesu110
Sesuvium_crithmoides_sesu83
Sesuvium_crithmoides_sesu108
Sesuvium_sp_cf_sesu81
Sesuvium_congense_sesu80
Sesuvium_sesuvioides_sesu128
Sesuvium_sesuvioides_gen_lmu
Sesuvium_sesuvioides_sesu134
Sesuvium_sesuvi_sesu120
Sesuvium_hydaspicum_sesu127
Sesuvium_hydaspicum_sesu104
Sesuvium_hydaspicum_sesu124
Sesuvium_nyasicum_sesu119
Sesuvium_nyasicum_sesu136
Sesuvium_hydaspicum_sesu13
Sesuvium_nyasicum_sesu111
Sesuvium_nyasicum_sesu138
Sesuvium_sp_nov_sesu130
Sesuvium_verrucosum_sesu4A
Sesuvium_verrucosum_trp
Sesuvium_sessile_sesu61
Sesuvium_portulacastrum_sesu74
Sesuvium_portulacastrum_trp
Sesuvium_humifusum_trp
Sesuvium_humifusum_sesu75
Zaleya_pentandra_trp
Zaleya_pentandra_sesu40
Zaleya_redimita_sesu95
Trianthema_corymbosa_sesu47
Trianthema_transvaalensis_sesu50
Trianthema_sheilae_sesu51
Trianthema_portulacastrum_trp
Trianthema_portulacastrum_trp2
Trianthema_ceratosepala_sesu73
Tribulocarpus_dimorphanthus_sesu142
Tribulocarpus_retusus_sesu72
Mesembryanthemum_crystallinum_trp
Delosperma_echinatum_trp
Beta_vulgaris_gen
Spinacia_oleracea_gen
Dianthus_caryophyllus_gen
Stegnosperma_halimifolium_trp

Fig. S3.

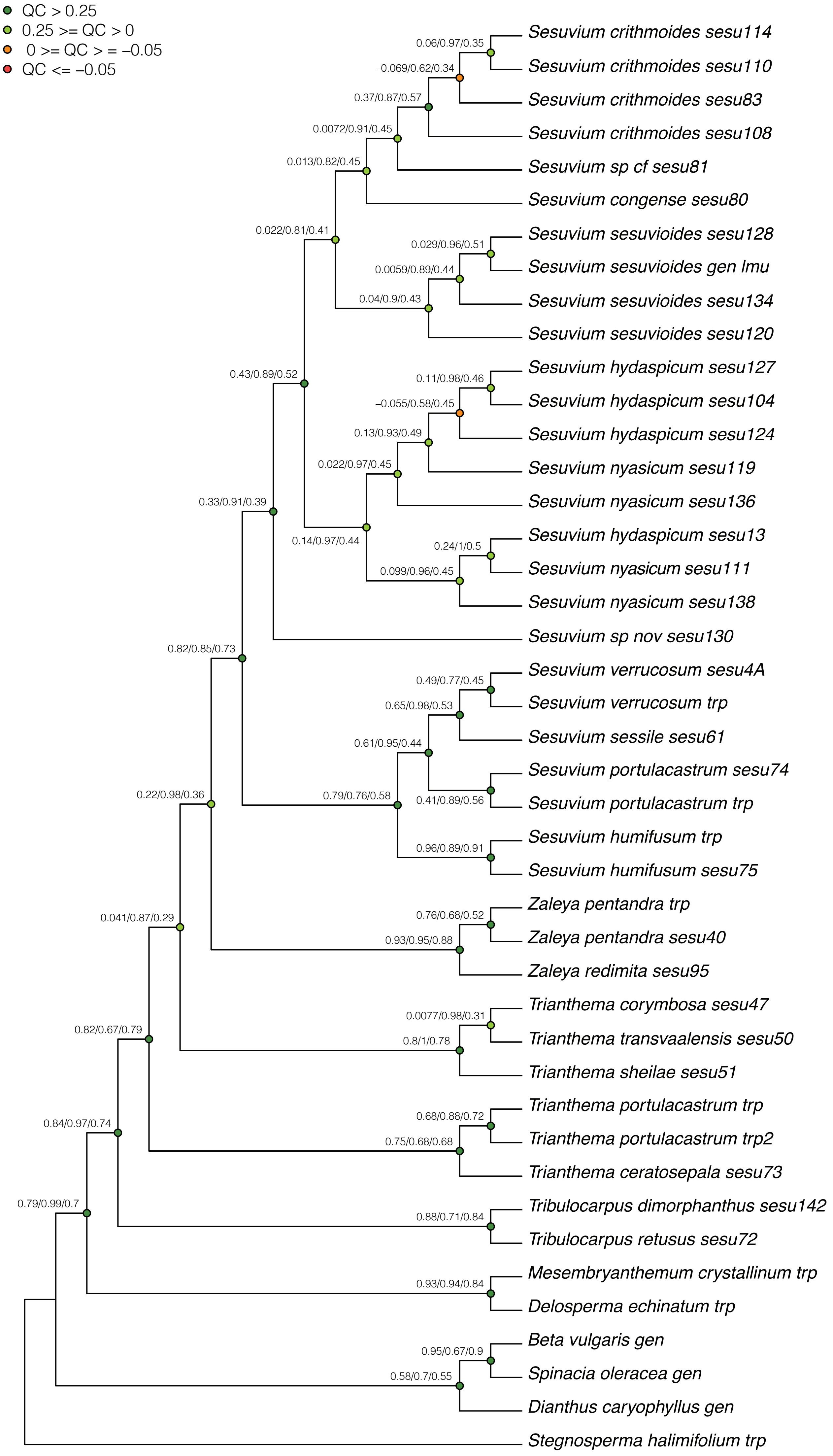

QC > 0.25
0.25 >= QC > 0
0 >= QC > = −0.05
QC <= −0.05
Sesuvium crithmoides sesu114
Sesuvium crithmoides sesu110
Sesuvium crithmoides sesu83
Sesuvium crithmoides sesu108
Sesuvium sp cf sesu81
Sesuvium congense sesu80
Sesuvium sesuvioides sesu128
Sesuvium sesuvioides gen lmu
Sesuvium sesuvioides sesu134
Sesuvium sesuvioides sesu120
Sesuvium hydaspicum sesu127
Sesuvium hydaspicum sesu104
Sesuvium hydaspicum sesu124
Sesuvium nyasicum sesu119
Sesuvium nyasicum sesu136
Sesuvium hydaspicum sesu13
Sesuvium nyasicum sesu111
Sesuvium nyasicum sesu138
Sesuvium sp nov sesu130
Sesuvium verrucosum sesu4A
Sesuvium verrucosum trp
Sesuvium sessile sesu61
Sesuvium portulacastrum sesu74
Sesuvium portulacastrum trp
Sesuvium humifusum trp
Sesuvium humifusum sesu75
Zaleya pentandra trp
Zaleya pentandra sesu40
Zaleya redimita sesu95
Trianthema corymbosa sesu47
Trianthema transvaalensis sesu50
Trianthema sheilae sesu51
Trianthema portulacastrum trp
Trianthema portulacastrum trp2
Trianthema ceratosepala sesu73
Tribulocarpus dimorphanthus sesu142
Tribulocarpus retusus sesu72
Mesembryanthemum crystallinum trp
Delosperma echinatum trp
Beta vulgaris gen
Spinacia oleracea gen
Dianthus caryophyllus gen
Stegnosperma halimifolium trp
0.06/0.97/0.35
−0.069/0.62/0.34
0.37/0.87/0.57
0.0072/0.91/0.45
0.013/0.82/0.45
0.022/0.81/0.41
0.029/0.96/0.51
0.0059/0.89/0.44
0.04/0.9/0.43
0.43/0.89/0.52
0.11/0.98/0.46
−0.055/0.58/0.45
0.13/0.93/0.49
0.022/0.97/0.45
0.33/0.91/0.39
0.14/0.97/0.44
0.24/1/0.5
0.099/0.96/0.45
0.82/0.85/0.73
0.49/0.77/0.45
0.65/0.98/0.53
0.61/0.95/0.44
0.22/0.98/0.36
0.79/0.76/0.58
0.41/0.89/0.56
0.96/0.89/0.91
0.041/0.87/0.29
0.76/0.68/0.52
0.93/0.95/0.88
0.0077/0.98/0.31
0.82/0.67/0.79
0.8/1/0.78
0.84/0.97/0.74
0.68/0.88/0.72
0.75/0.68/0.68
0.79/0.99/0.7
0.88/0.71/0.84
0.93/0.94/0.84
0.95/0.67/0.9
0.58/0.7/0.55

Fig. S4.

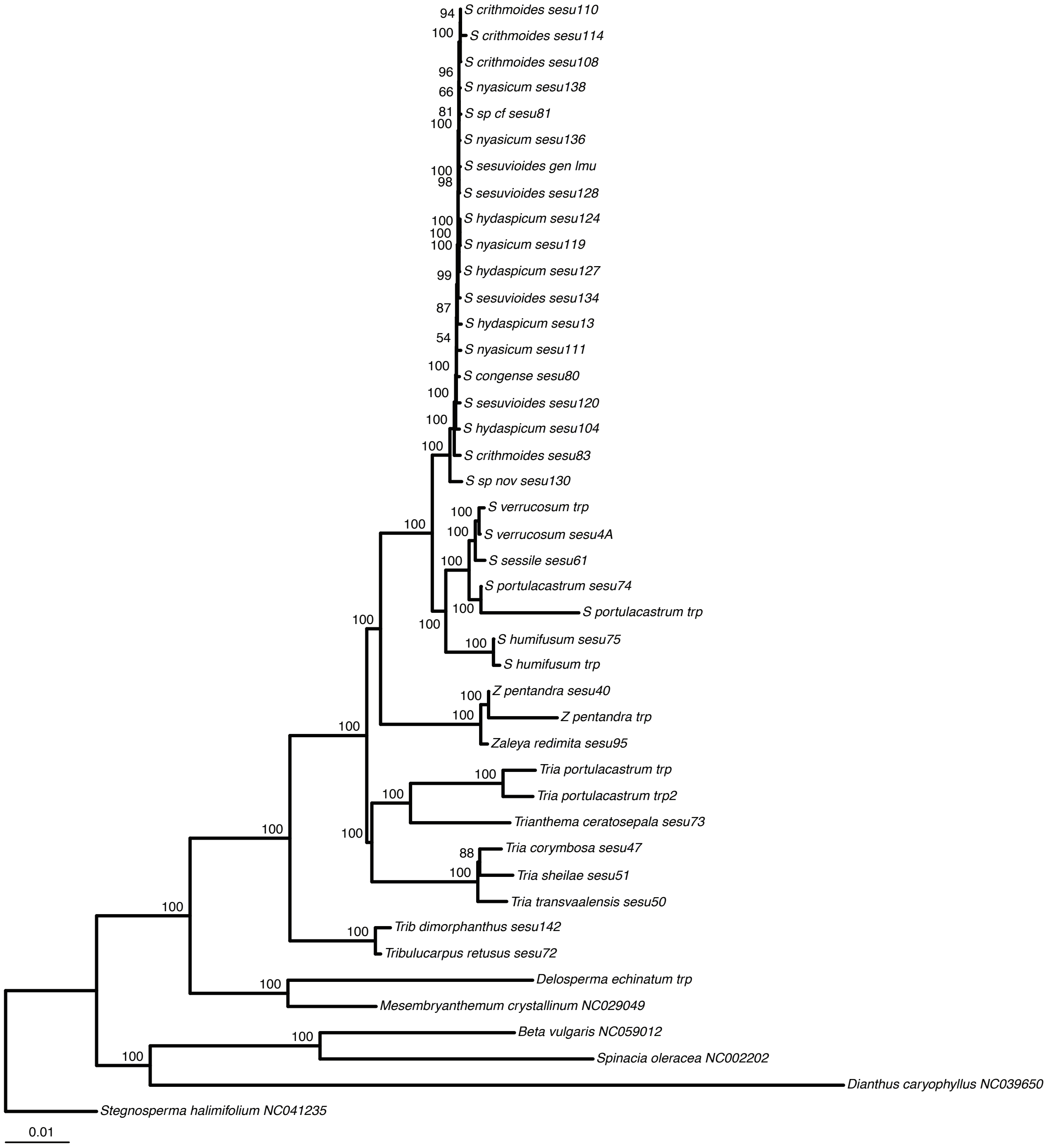
S crithmoides sesu110
S crithmoides sesu114
S crithmoides sesu108
S nyasicum sesu138
S sp cf sesu81
S nyasicum sesu136
S sesuvioides gen lmu
S sesuvioides sesu128
S hydaspicum sesu124
S nyasicum sesu119
S hydaspicum sesu127
S sesuvioides sesu134
S hydaspicum sesu13
S nyasicum sesu111
S congense sesu80
S sesuvioides sesu120
S hydaspicum sesu104
S crithmoides sesu83
S sp nov sesu130
S verrucosum trp
S verrucosum sesu4A
S sessile sesu61
S portulacastrum sesu74
S portulacastrum trp
S humifusum sesu75
S humifusum trp
Z pentandra sesu40
Z pentandra trp
Zaleya redimita sesu95
Tria portulacastrum trp
Tria portulacastrum trp2
Trianthema ceratosepala sesu73
Tria corymbosa sesu47
Tria sheilae sesu51
Tria transvaalensis sesu50
Trib dimorphanthus sesu142
Tribulucarpus retusus sesu72
Delosperma echinatum trp
Mesembryanthemum crystallinum NC029049
Beta vulgaris NC059012
Spinacia oleracea NC002202
Dianthus caryophyllus NC039650
Stegnosperma halimifolium NC041235
0.01

Fig. S5.

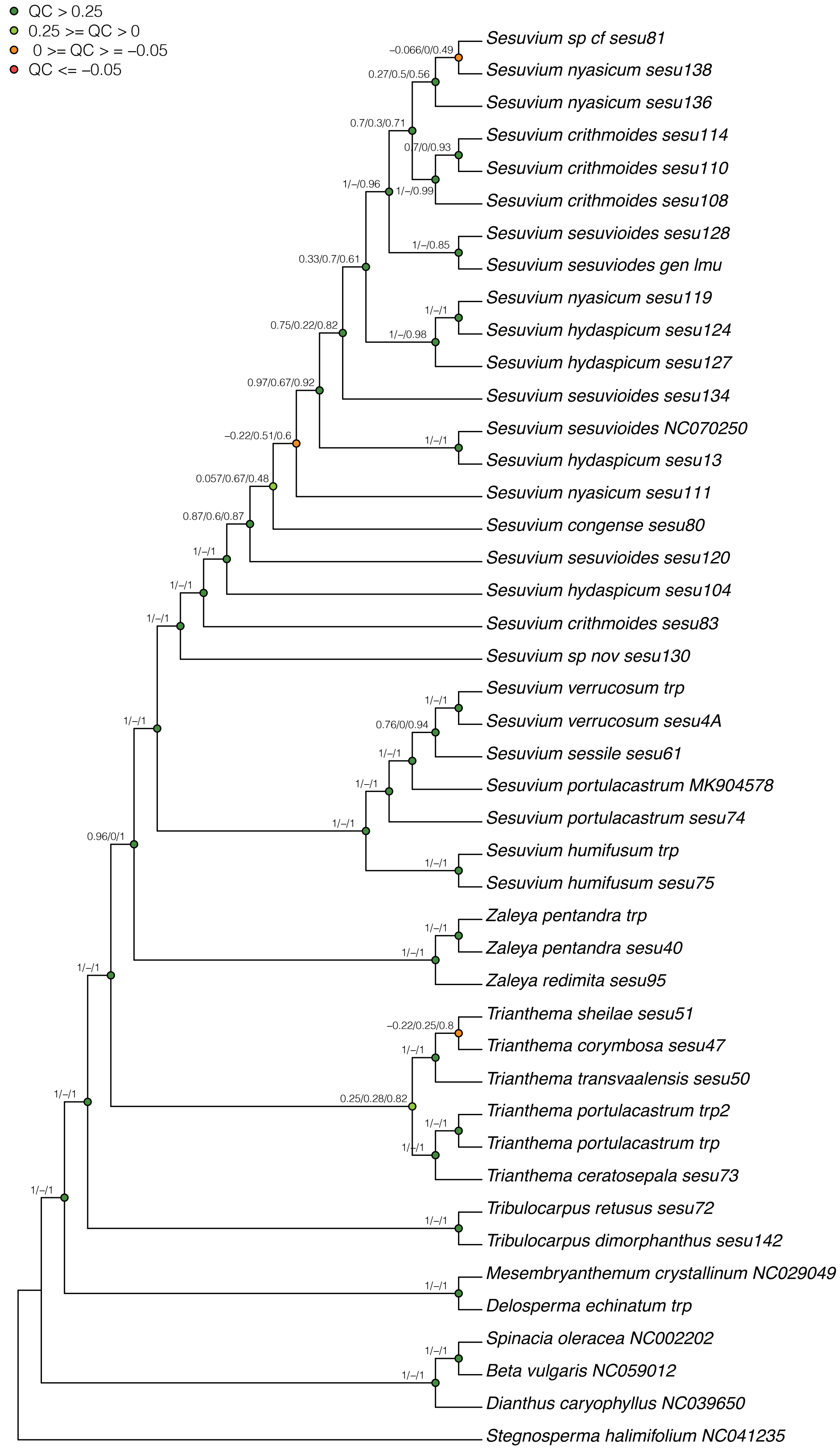
QC > 0.25
0.25 >= QC > 0
0 >= QC > = −0.05
QC <= −0.05
Sesuvium sp cf sesu81
Sesuvium nyasicum sesu138
Sesuvium nyasicum sesu136
Sesuvium crithmoides sesu114
Sesuvium crithmoides sesu110
Sesuvium crithmoides sesu108
Sesuvium sesuvioides sesu128
Sesuvium sesuviodes gen lmu
Sesuvium nyasicum sesu119
Sesuvium hydaspicum sesu124
Sesuvium hydaspicum sesu127
Sesuvium sesuvioides sesu134
Sesuvium sesuvioides NC070250
Sesuvium hydaspicum sesu13
Sesuvium nyasicum sesu111
Sesuvium congense sesu80
Sesuvium sesuvioides sesu120
Sesuvium hydaspicum sesu104
Sesuvium crithmoides sesu83
Sesuvium sp nov sesu130
Sesuvium verrucosum trp
Sesuvium verrucosum sesu4A
Sesuvium sessile sesu61
Sesuvium portulacastrum MK904578
Sesuvium portulacastrum sesu74
Sesuvium humifusum trp
Sesuvium humifusum sesu75
Zaleya pentandra trp
Zaleya pentandra sesu40
Zaleya redimita sesu95
Trianthema sheilae sesu51
Trianthema corymbosa sesu47
Trianthema transvaalensis sesu50
Trianthema portulacastrum trp2
Trianthema portulacastrum trp
Trianthema ceratosepala sesu73
Tribulocarpus retusus sesu72
Tribulocarpus dimorphanthus sesu142
Mesembryanthemum crystallinum NC029049
Delosperma echinatum trp
Spinacia oleracea NC002202
Beta vulgaris NC059012
Dianthus caryophyllus NC039650
Stegnosperma halimifolium NC041235
−0.066/0/0.49
0.27/0.5/0.56
0.7/0.3/0.71
0.7/0/0.93
1/–/0.96
1/–/0.99
1/–/0.85
0.33/0.7/0.61
1/–/1
0.75/0.22/0.82
1/–/0.98
0.97/0.67/0.92
−0.22/0.51/0.6
0.057/0.67/0.48
0.87/0.6/0.87
0.76/0/0.94
0.96/0/1
−0.22/0.25/0.8
0.25/0.28/0.82

Fig. S6.

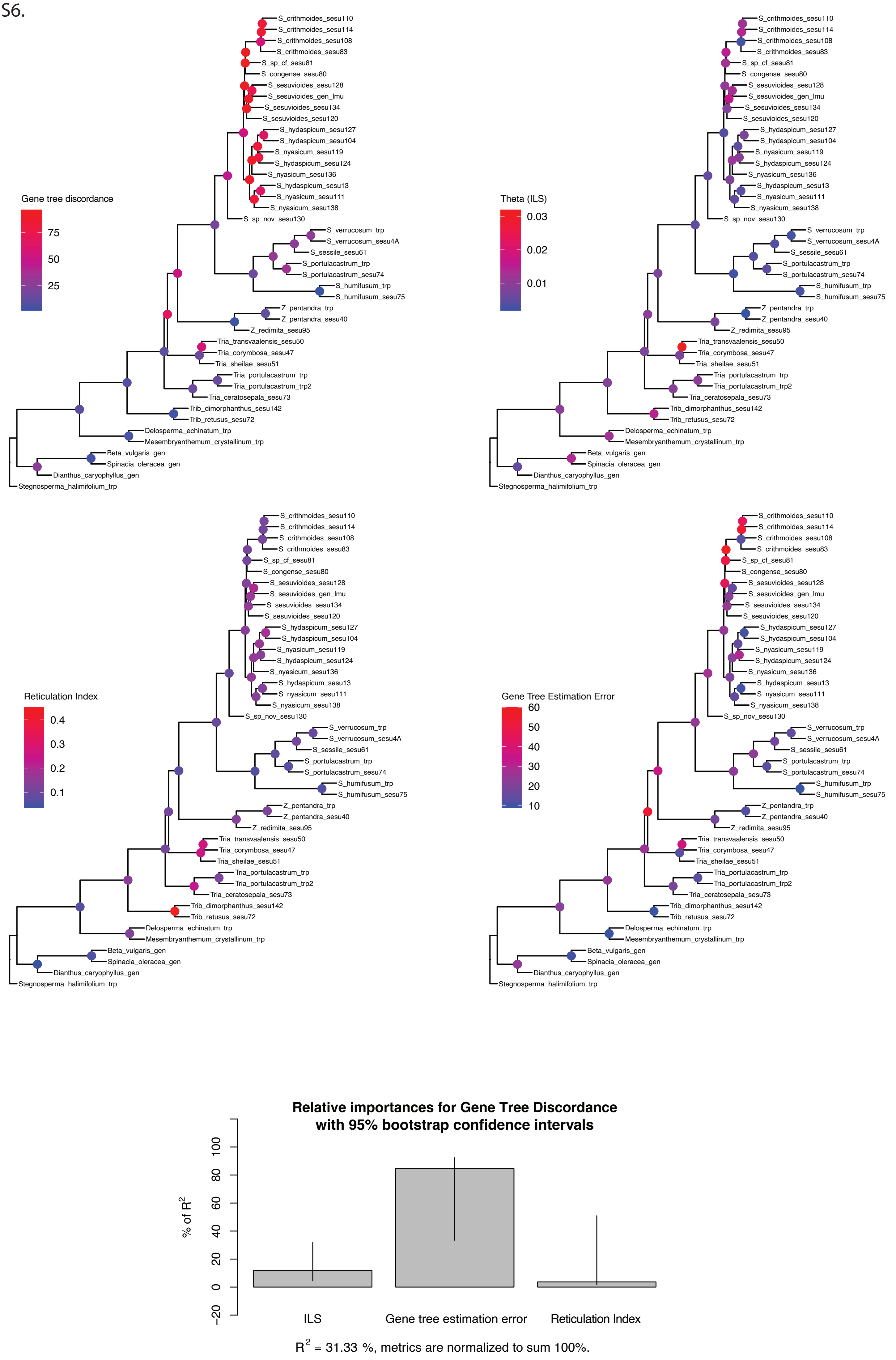
Gene tree discordance
75
50
25
Theta (ILS)
0.03
0.02
0.01
Reticulation Index
0.4
0.3
0.2
0.1
Gene Tree Estimation Error
60
50
40
30
20
10
S_crithmoides_sesu110
S_crithmoides_sesu114
S_crithmoides_sesu108
S_crithmoides_sesu83
S_sp_cf_sesu81
S_congense_sesu80
S_sesuvioides_sesu128
S_sesuvioides_gen_lmu
S_sesuvioides_sesu134
S_sesuvioides_sesu120
S_hydaspicum_sesu127
S_hydaspicum_sesu104
S_nyasicum_sesu119
S_hydaspicum_sesu124
S_nyasicum_sesu136
S_hydaspicum_sesu13
S_nyasicum_sesu111
S_nyasicum_sesu138
S_sp_nov_sesu130
S_verrucosum_trp
S_verrucosum_sesu4A
S_sessile_sesu61
S_portulacastrum_trp
S_portulacastrum_sesu74
S_humifusum_trp
S_humifusum_sesu75
Z_pentandra_trp
Z_pentandra_sesu40
Z_redimita_sesu95
Tria_transvaalensis_sesu50
Tria_corymbosa_sesu47
Tria_sheilae_sesu51
Tria_portulacastrum_trp
Tria_portulacastrum_trp2
Tria_ceratosepala_sesu73
Trib_dimorphanthus_sesu142
Trib_retusus_sesu72
Delosperma_echinatum_trp
Mesembryanthemum_crystallinum_trp
Beta_vulgaris_gen
Spinacia_oleracea_gen
Dianthus_caryophyllus_gen
Stegnosperma_halimifolium_trp
Relative importances for Gene Tree Discordance
with 95% bootstrap confidence intervals
% of R2
100
80
60
40
20
0
−20
ILS
Gene tree estimation error
Reticulation Index
R2 = 31.33 %, metrics are normalized to sum 100%.

Fig. S7.

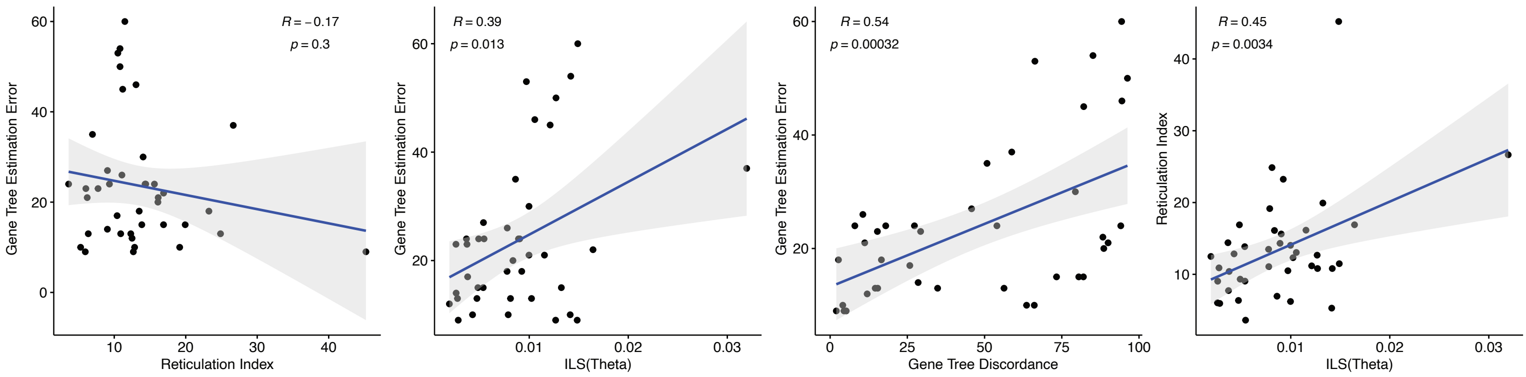
R = −0.17
p = 0.3
Gene Tree Estimation Error
Reticulation Index
R = 0.39
p = 0.013
Gene Tree Estimation Error
ILS(Theta)
R = 0.54
p = 0.00032
Gene Tree Estimation Error
Gene Tree Discordance
R = 0.45
p = 0.0034
Reticulation Index
ILS(Theta)

Fig. S8.

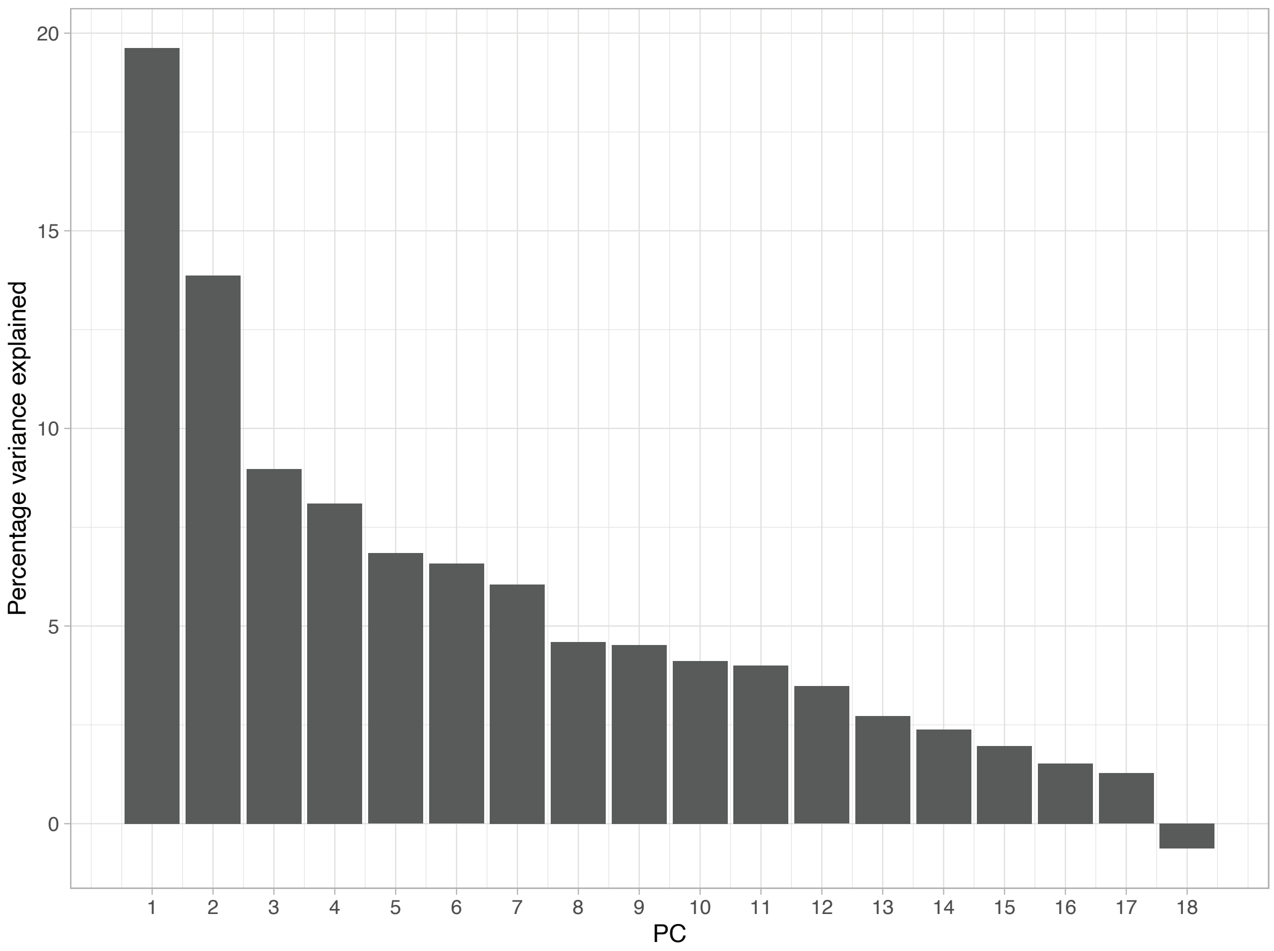
Percentage variance explained
20
15
10
5
0
1 2 3 4 5 6 7 8 9 10 11 12 13 14 15 16 17 18
PC

Fig. S9.

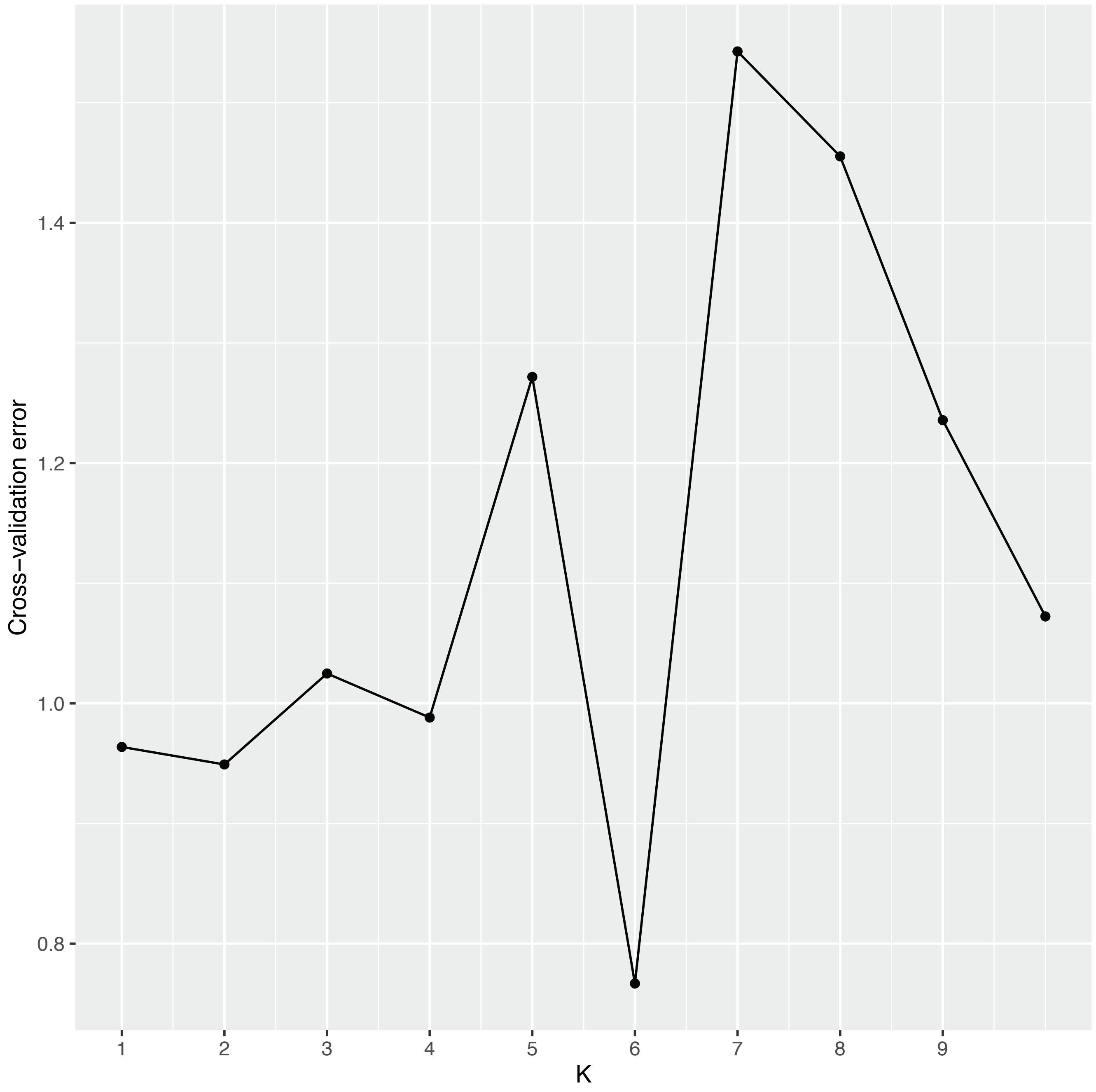

Fig. S10.

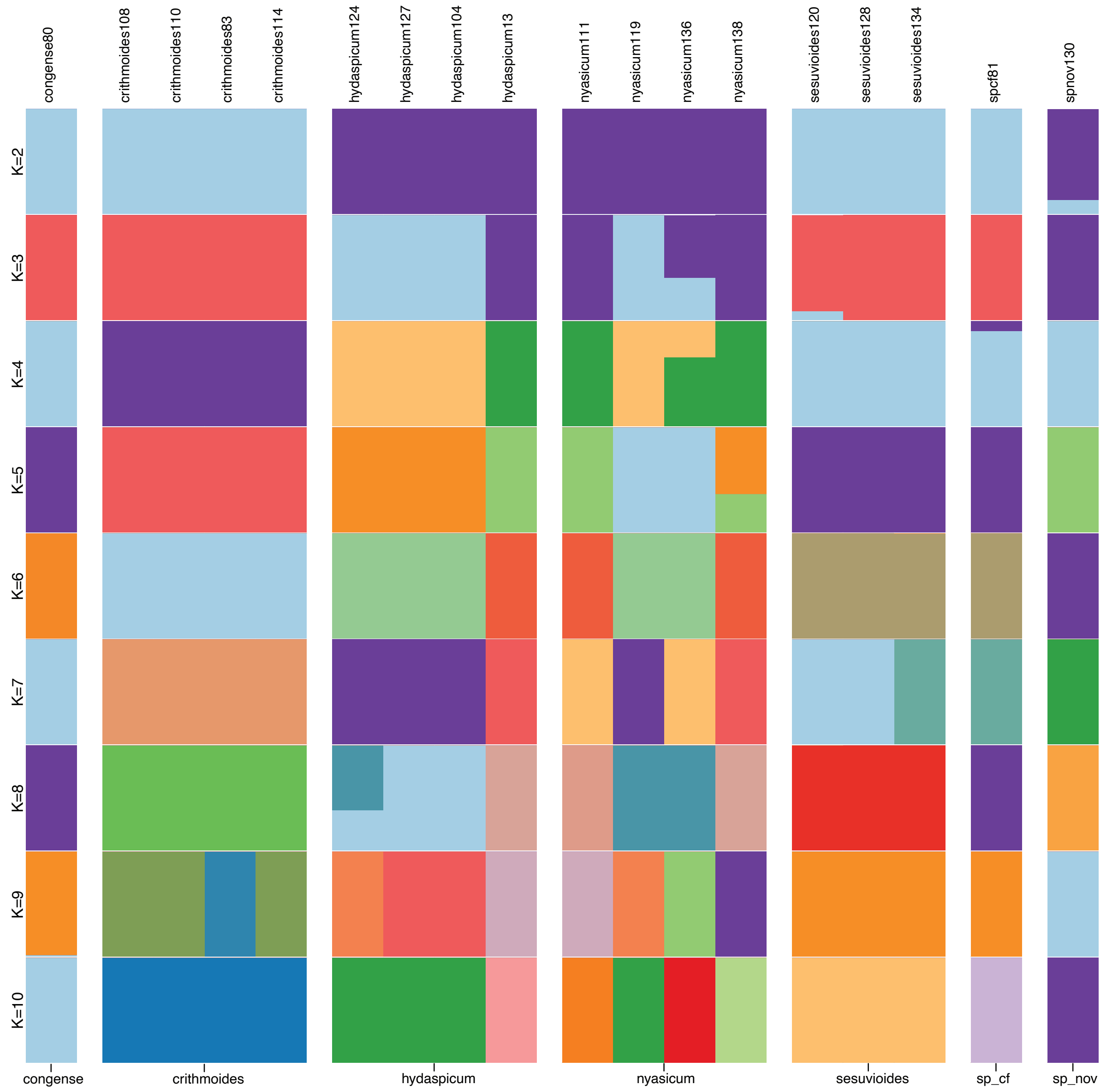
congense80
crithmoides108
crithmoides110
crithmoides83
crithmoides114
hydaspicum124
hydaspicum127
hydaspicum104
hydaspicum13
nyasicum111
nyasicum119
nyasicum136
nyasicum138
sesuvioides120
sesuvioides128
sesuvioides134
spcf81
spnov130
K=2
K=3
K=4
K=5
K=6
K=7
K=8
K=9
K=10
congense
crithmoides
hydaspicum
nyasicum
sesuvioides
sp_cf
sp_nov

Fig. S11.

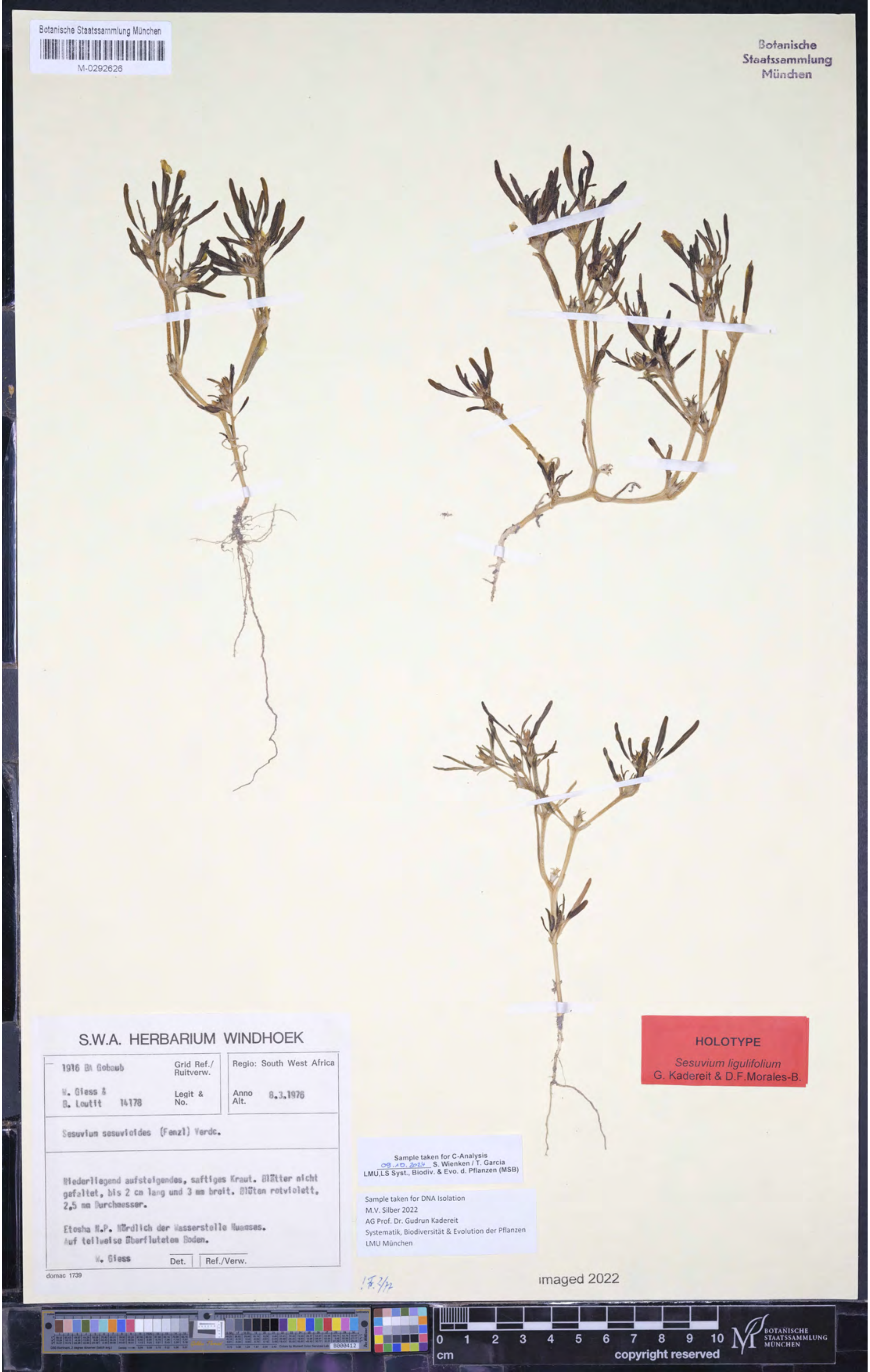
Botanische Staatssammlung München
M-0292626
Botanische Staatssammlung München
S.W.A. HERBARIUM WINDHOEK
1916 BA Gobaub
Grid Ref./ Ruitverw.
Regio: South West Africa
W. Giess & B. Loutit 14178
Legit & No.
Anno Alt. 8.3.1976
Sesuvium sesuvioides (Fenzl) Verdc.
Niederliegend aufsteigendes, saftiges Kraut. Blätter nicht gefaltet, bis 2 cm lang und 3 mm breit. Blüten rotviolett, 2,5 mm Durchmesser.
Etosha N.P. Nördlich der Wasserstelle Nuamses. Auf teilweise überflutetem Boden.
W. Giess
Det.
Ref./Verw.
domac 1739
HOLOTYPE
Sesuvium ligulifolium
G. Kadereit & D.F.Morales-B.
Sample taken for C-Analysis
09.10.2024 S. Wienken / T. Garcia
LMU,LS Syst., Biodiv. & Evo. d. Pflanzen (MSB)
Sample taken for DNA Isolation
M.V. Silber 2022
AG Prof. Dr. Gudrun Kadereit
Systematik, Biodiversität & Evolution der Pflanzen
LMU München
imaged 2022
0 1 2 3 4 5 6 7 8 9 10
cm
copyright reserved
BOTANISCHE STAATSSAMMLUNG MÜNCHEN

Table S1. Taxon sampling and voucher information for newly sequenced samples.

| Lib ID | Taxon | DNA ID | Country | Collected by | Specimen Voucher | Collection date | Origin | SRA | GenBank |
|---|---|---|---|---|---|---|---|---|---|
| S11 | *Sesuvium congense* | sesu80 | Angola | P. J. D. Winter 7779 | PRE 849008.0 | 23.01.2009 | N of Namibe. Road to Baba from Lucira rd. | | |
| S04 | *Sesuvium crithmoides* | sesu108 | Congo | Gillet J. s.n. | BR 0000013827434 | n.a. | Kongo Central prov.: Banana, Congo prés Gillet | | |
| S13 | *Sesuvium crithmoides* | sesu83 | Angola | P. J. D. Winter 7786 | PRE 849042.0 | 23.01.2009 | Baba. Coast W of Namibe-Lucira Rd. | | |
| S35 | *Sesuvium crithmoides* | sesu110 | Congo | Flamigni A. 10773 | BR 0000013827427 | 1959 | Kongo Central prov.: Nature Reserve Luki-Mayumbe | | |
| S24 | *Sesuvium crithmoides* | sesu114 | Angola | Lebrun J. 10905 | BR 0000013827403 | 13.09.1955 | Luanda prov.: Luanda | | |
| S18 | *Sesuvium hydaspicum* | sesu124 | Namibia | Bertil Nordenstam 2305 | M-0292641 | 20.04.1963 | Maltahöhe Distr.: 26 miles W. of Maltahöhe on Naukluft road, sandy | | |
| S25 | *Sesuvium hydaspicum* | sesu127 | Namibia | B. de Winter 3577 | M-0292644 | 17.05.1955 | 9.9 m. from Ariamsvlei on rd. to Charlies Puts | | |
| S01 | *Sesuvium hydaspicum* | sesu13 | Burkina Faso | J.E. Madsen 5771 | S05-5896 | 21.09.1996 | Mare de Menegou, Oudalan, large swamp with aquatic plants and | | |
| S23 | *Sesuvium hydaspicum* | sesu104 | Namibia | A. Sukhorukov 65 | MW 0582873 | 07.03.2017 | Karas Region, Goargeb | | |

| | | | | | | | |
|---|---|---|---|---|---|---|---|
| S15 | *Sesuvium nyasicum* | sesu111 | Zimbabwe | Symoens J. 13910 | BR 0000013829 124 | 25.12.1970 | n.a. |
| S16 | *Sesuvium nyasicum* | sesu119 | Namibia | P.M.Burgoyne, N.Snow 5074 | MO 5667010 | 03.01.1996 | Rehoboth, 4km von Rehoboth |
| S27 | *Sesuvium nyasicum* | sesu136 | Namibia | B. de Winter & Leistner 5258 | M-0292633 | 29.03.1963 | Kaokoveld |
| S28 | *Sesuvium nyasicum* | sesu138 | Namibia | Robert J. Rodin 9392 | M-0292635 | 04.04.1973 | Just west of Oshikango |
| S21 | *Sesuvium portulacastrum* | sesu74 | Mexico | Sage s. n. | MJG 014143 | Sep-2010 | Yucatan (cultivated at Univ. Toronto) |
| S10 | *Sesuvium sessile* | sesu61 | Mexico | Mora-Olivo 7026 | MEXU 1231179 | 10.11.08 | Tamaulipas, 4 km N of Poblado Anáhuac, Mpio. Valle Hermoso |
| S12 | *Sesuvium spec.* | sesu81 | Angola | P. J. D. Winter 7677 | PRE 848975.0 | 18.01.2009 | Namibe, 25 km W of Caraculo |
| S17 | *Sesuvium sesuvioides* | sesu120 | Namibia | Bertil Nordenstam 2491 | M-0292639 | 03.05.1963 | Omaruru Distr.: Brandberg, Tsisab Valley, along the |
| S29 | *Sesuvium sesuvioides* | sesu134 | Namibia | Ihlenfeldt, de Winter & Hardy 3137 | M-0292630 | 24.03.1963 | Swartbankberge am Kuiseb |
| S30 | *Sesuvium sesuvioides* | sesu128 | Namibia | W. Giess & M. Müller 14452 | M-0292632 | 17.06.1976 | Fars Altdorn |
| S26 | *Sesuvium spec. nov.* | sesu130 | Namibia | W. Giess & B. Loutit 14178 | M-0292626 | 03.08.1975 | Etosha National park, South West Africa |

| | | | | | | | |
|---|---|---|---|---|---|---|---|
| S31 | *Sesuvium verrucosum* | sesu4A | USA | Steve Boyd et al. 3549 | RSA 0325961 | 07.05.1989 | California, Riverside, Vail Lake area, pond and freshwater marsh in |
| S22 | *Sesuvium humifusum* | sesu75 | USA | Sage s. n. | MJG 014141 | n.a. | Nevada, Spangler Springs Reservoir |
| S03 | *Trianthema ceratosepala* | sesu73 | Somalia | Thulin, Hedrén & Dahir 7332 | UPS (V-51249) 3029 | 08.05.1990 | Galguduud, 18km on road from Ceelbuur to Ceeldheer |
| S33 | *Trianthema corymbosa* | sesu47 | South Africa | Hartmann & Potgieter 32657 | HBG s. n. | 11.03.1995 | Northern Cape, Namaqualand, Numees |
| S02 | *Trianthema sheilae* | sesu51 | Sudan | Hartmann 21470 | HBG s. n. | 17.11.1987 | 81 mls N of Aroma |
| S34 | *Trianthema transvaalensis* | sesu50 | South Africa | Hartmann & Dehn 25119 | HBG s. n. | 09.02.1988 | Natal/KwaZulu/Ubombo, Mzuke Park, between gate and officebuilding |
| S36 | *Tribulocarpus dimorphanthus* | sesu142 | n.a. | Ihlenfeldt 3040 | HBG s. n. | | n.a. |
| S20 | *Tribulocarpus retusus* | sesu72 | Somalia | Thulin, Dahir, Khalid & Osman 10511 | UPS V-113019 | 09.05.2011 | Nugaal region: 19 km along track from Gaalogod to Garadeen |
| S09 | *Zaleya pentandra* | sesu40 | Saudi Arabia | I. & O. Hedberg 92042 | UPS (V-60916) 66393 | 07.05.1992 | Jizan, Fayfa, around the top station |
| S14 | *Zaleya redimita* | sesu95 | Kenya | HEK 21408 | HBG s. n. | 15.03.1987 | Eastern Province: ca. 40 km W Marsabit nach Furole |

Table S2. Transcriptome and genome used for tree-based othololgous loci inference.

| **Sample** | **Family** | **Type** | **Original raw data source** | **Assembly and translation (CDS) source** |
|---|---|---|---|---|
| *Delosperma echinatum* | Aizoaceae | Transcriptome | Matasci et al., 2014 | Yang et al., 2015 |
| *Mesembryanthemum crystallinum* | Aizoaceae | Transcriptome | Christin et al., 2015 | Yang et al., 2015 |
| *Sesuvium humifusum* | Aizoaceae | Transcriptome | Matasci et al., 2014 | Yang et al., 2015 |
| *Sesuvium portulacastrum* | Aizoaceae | Transcriptome | Matasci et al., 2014 | Yang et al., 2015 |
| *Sesuvium verrucosum* | Aizoaceae | Transcriptome | Matasci et al., 2014 | Yang et al., 2015 |
| *Trianthema portulacastrum (trp1)* | Aizoaceae | Transcriptome | Matasci et al., 2014 | Yang et al., 2015 |
| *Trianthema portulacastrum (trp2)* | Aizoaceae | Transcriptome | Christin et al., 2015 | Yang et al., 2015 |
| *Zaleya pentandra* | Aizoaceae | Transcriptome | Matasci et al., 2014 | Yang et al., 2015 |
| *Beta vulgaris* | Amaranthaceae | Genome | Dohm et al., 2014 | Yang et al., 2015 |
| *Dianthus caryophyllus* | Caryophyllaceae | Genome | Yagi et al., 2014 | Yang et al., 2015 |
| *Spinacia oleracea* | Amaranthaceae | Genome | Dohm et al., 2014 | Yang et al., 2015 |
| *Stegnosperma halimifolium* | Stegnospermatacea | Transcriptome | Yang et al., 2018 | Yang et al., 2018 |

Table S3. Additional samples used for plastome assembly and phylogenetic analyses.

| Sample | Family | Type | SRA accession | GenBank accession | Usage | Source |
|---|---|---|---|---|---|---|
| *Sesuvium humifusum* | Aizoaceae | Transcriptome | ERR2040193 | N/A | Phylogenetic analyses | Matasci et al., 2014 |
| *Delosperma echinatum* | Aizoaceae | Transcriptome | ERR2040192 | N/A | Phylogenetic analyses | Matasci et al., 2014 |
| *Mesembryanthemum crystallinum* | Aizoaceae | Plastome | N/A | NC029049 | Phylogenetic analyses | GenBank - unpublished |
| *Sesuvium portulacastrum* | Aizoaceae | Plastome | N/A | MK904578 | Reference for assembly | GenBank- unpublished |
| *Sesuvium portulacastrum* | Aizoaceae | Transcriptome | ERR2040194 | N/A | Phylogenetic analyses | Matasci et al., 2014 |
| *Sesuvium verrucosum* | Aizoaceae | Transcriptome | ERR2040195+ ERR2040196 | N/A | Phylogenetic analyses | Matasci et al., 2014 |
| *Sesuvium sesuviodes* | Aizoaceae | Plastome | N/A | NC070250 | Reference for assembly | Javaid et al., 2023 |
| *Sesuvium sesuviodes* | Aizoaceae | Plastome | N/A | N/A | Phylogenetic analyses | Siadjeu and Kadereit (unpublished) |
| *Tetragonia tetragonioides* | Aizoaceae | Plastome | N/A | NC036991 | Reference for assembly | Choi et al., 2018 |
| *Trianthema portulacastrum* | Aizoaceae | Transcriptome | ERR2040197+ ERR2040198 | N/A | Phylogenetic analyses | Matasci et al., 2014 |
| *Trianthema portulacastrum* | Aizoaceae | Transcriptome | SRR1698227+ SRR1698228 | N/A | Phylogenetic analyses | Christin et al., 2015 |
| *Zaleya_pentandra* | Aizoaceae | Plastome | N/A | MN296416 | Reference for assembly | Xu, 2019 |
| *Zaleya pentandra* | Aizoaceae | Transcriptome | ERR2040199 | N/A | Phylogenetic analyses | Matasci et al., 2014 |
| *Beta vulgaris* | Amaranthaceae | Plastome | N/A | NC059012 | Phylogenetic analyses | Sielemann et al., 2022 |
| *Dianthus caryophyllus* | Caryophyllaceae | Plastome | N/A | NC039650 | Phylogenetic analyses | Chen et al., 2018 |
| *Spinacia oleracea* | Amaranthaceae | Plastome | N/A | NC002202 | Phylogenetic analyses | Schmitz-Linneweber et al., 2011 |

| | | | | | | |
|---|---|---|---|---|---|---|
| *Stegnosperma halimifolium* | Stegnospermataceae | Plastome | N/A | NC041235 | Phylogenetic analyses | Yao et al., 2019 |

Table S4. Sequencing, orthologs, paralogs, and plastome assembly statistics

| Sample - final names (used in Figures and main text) | Raw reads (million) | After deduplication reads (million) | Clean reads (million) | Genes recovered | Percentage of genes (from 9,151) | Paralogs | Orthologs | Percentage of orthologs (from 7,138) | Number of aligned bases | Percentage of aligned bases (from 10,461,324) | Plastome assembly | Plastome length | Plastome length (without IRA) |
|---|---|---|---|---|---|---|---|---|---|---|---|---|---|
| Sesuvium_congense_sesu80 | 28.63 | 24.86 | 24.52 | 9,123 | 99.7 | 2,577 | 6844 | 95.88 | 9215497 | 88.09 | Full | 155976 | 130207 |
| Sesuvium_crithmoides_sesu110 | 24.95 | 20.34 | 20.11 | 8,363 | 91.4 | 1,263 | 6015 | 84.27 | 5863784 | 56.05 | Full | 155964 | 130200 |
| Sesuvium_crithmoides_sesu83 | 25.96 | 21.3 | 21.06 | 9,110 | 99.6 | 2,327 | 6848 | 95.94 | 9238181 | 88.31 | Full | 155986 | 130217 |
| Sesuvium_sesuvioides_sesu120 | 22.17 | 21.08 | 20.85 | 6,891 | 75.3 | 409 | 4765 | 66.76 | 3544410 | 33.88 | Full | 155908 | 130144 |
| Sesuvium_hydaspicum_sesu124 | 29.86 | 24.37 | 24.12 | 9,128 | 99.7 | 2,855 | 6930 | 97.09 | 9446587 | 90.3 | Full | 155895 | 130126 |
| Sesuvium_hydaspicum_sesu127 | 22.57 | 16.31 | 16.05 | 9,123 | 99.7 | 2,676 | 6895 | 96.6 | 9610988 | 91.87 | Full | 155892 | 130123 |
| Sesuvium_hydaspicum_sesu104 | 20.38 | 15.1 | 14.93 | 9,123 | 99.7 | 2,450 | 6880 | 96.39 | 9479941 | 90.62 | Full | 155958 | 130179 |
| Sesuvium_hydaspicum_sesu111 | 26.64 | 24.58 | 24.19 | 8,687 | 94.9 | 873 | 6326 | 88.62 | 6357920 | 60.78 | Full | 155936 | 130168 |
| Sesuvium_nyasicum_sesu119 | 23.18 | 20.39 | 20.22 | 8,815 | 96.3 | 1,391 | 6489 | 90.91 | 7037660 | 67.27 | Full | 155907 | 130138 |
| Sesuvium_nyasicum_sesu136 | 28.53 | 21.99 | 21.74 | 9,108 | 99.5 | 2,674 | 6886 | 96.47 | 9179291 | 87.75 | Full | 155964 | 130194 |

| | | | | | | | | | | | | | |
|---|---|---|---|---|---|---|---|---|---|---|---|---|---|
| Sesuvium_nyasicum_sesu138 | 20.79 | 15.77 | 15.59 | 9,114 | 99.6 | 2,464 | 6909 | 96.79 | 9305114 | 88.95 | Full | 155930 | 130160 |
| Sesuvium_portulacastrum_sesu74 | 21.42 | 15.27 | 15.04 | 9,138 | 99.9 | 6,000 | 6880 | 96.39 | 8911523 | 85.19 | Full | 156187 | 130418 |
| Sesuvium_sesuvioides_sesu128 | 22.32 | 17.78 | 17.57 | 9,105 | 99.5 | 2,533 | 6800 | 95.26 | 8882245 | 84.91 | Full | 155879 | 130110 |
| Sesuvium_sesuvioides_sesu134 | 26.79 | 22.5 | 22.23 | 8,872 | 97 | 1,573 | 6534 | 91.54 | 7373723 | 70.49 | Full | 155895 | 130125 |
| Sesuvium_sp_cf_sesu81 | 19.81 | 15.77 | 15.59 | 8,812 | 96.3 | 1,562 | 6440 | 90.22 | 7079072 | 67.67 | Full | 155926 | 130150 |
| Sesuvium_verrucosum_sesu4A | 22.05 | 18.19 | 18.02 | 9,103 | 99.5 | 1,993 | 6591 | 92.34 | 8649693 | 82.68 | Full | 156040 | 130276 |
| Sesuvium_humifusum_sesu75 | 24.6 | 17.41 | 17.18 | 9,125 | 99.7 | 4,541 | 6545 | 91.69 | 8905705 | 85.13 | Full | 155546 | 129821 |
| Sesuvium_crithmoides_sesu108 | 34.6 | 33.87 | 33.58 | 4,657 | 50.9 | 204 | 2957 | 41.43 | 1734834 | 16.58 | Partial | --- | 129213 |
| Sesuvium_crithmoides_sesu114 | 34.6 | 25.24 | 24.9 | 8,787 | 96 | 1,828 | 6428 | 90.05 | 6932251 | 66.27 | Partial | --- | 129692 |
| Sesuvium_hydaspicum_sesu13 | 34.42 | 33.17 | 32.93 | 9,100 | 99.4 | 1,662 | 6829 | 95.67 | 9001882 | 86.05 | Partial | --- | 129710 |
| Sesuvium_sessile_sesu61 | 24.1 | 22.2 | 22.01 | 9,121 | 99.7 | 1,890 | 6567 | 92 | 8645449 | 82.64 | Partial | --- | 130444 |
| Sesuvium_sp_nov_sesu130 | 24.05 | 19.27 | 19.09 | 9,113 | 99.6 | 2,106 | 6876 | 96.33 | 9101057 | 87 | Partial | --- | 127302 |
| Trianthema_ceratosepala_sesu73 | 35.02 | 34.23 | 34.03 | 7,544 | 82.4 | 459 | 5533 | 77.51 | 4420190 | 42.25 | Partial | --- | 125650 |

| | | | | | | | | | | | | | |
|---|---|---|---|---|---|---|---|---|---|---|---|---|---|
| Trianthema_corymbosa_sesu47 | 22.97 | 18.19 | 18.01 | 8,970 | 98 | 2,358 | 6729 | 94.27 | 8090008 | 77.33 | Partial | --- | 127509 |
| Trianthema_sheilae_sesu51 | 26.73 | 24.97 | 24.4 | 3,239 | 35.4 | 191 | 2060 | 28.86 | 1084323 | 10.37 | Partial | --- | 127965 |
| Trianthema_transvaalensis_sesu50 | 22.23 | 17.32 | 17.14 | 8,967 | 98 | 2,680 | 6723 | 94.19 | 7739495 | 73.98 | Partial | --- | 127655 |
| Tribulocarpus_dimorphanthus_sesu142 | 23.57 | 20.33 | 20.14 | 7,534 | 82.3 | 821 | 5436 | 76.16 | 4451272 | 42.55 | Partial | --- | 125717 |
| Tribulocarpus_retusus_sesu72 | 24.3 | 17.22 | 16.99 | 9,130 | 99.8 | 2,672 | 6943 | 97.27 | 9366997 | 89.54 | Partial | --- | 130075 |
| Zaleya_pentandra_sesu40 | 26.9 | 22.5 | 22.29 | 9,130 | 99.8 | 2,385 | 7021 | 98.36 | 9496161 | 90.77 | Partial | --- | 129123 |
| Zaleya_redimita_sesu95 | 26.81 | 23.74 | 23.54 | 9,123 | 99.7 | 2,135 | 6995 | 98 | 9287958 | 88.78 | Partial | --- | 123009 |
| Beta_vulgaris_gen | --- | --- | --- | --- | --- | --- | 6551 | 91.78 | 8895286 | 85.03 | --- | --- | --- |
| Sesuvium_humifusum_trp | --- | --- | --- | --- | --- | --- | 5562 | 77.92 | 6394586 | 61.13 | Partial | | 64248 |
| Delosperma_echinatum_trp | --- | --- | --- | --- | --- | --- | 5442 | 76.24 | 4865092 | 46.51 | Partial | | 63460 |
| Dianthus_caryophyllus_gen | --- | --- | --- | --- | --- | --- | 4252 | 59.57 | 5744510 | 54.91 | --- | --- | --- |
| Mesembryanthemum_crystallinum_trp | --- | --- | --- | --- | --- | --- | 6556 | 91.85 | 7564592 | 72.31 | --- | --- | --- |
| Sesuvium_portulacastrum_trp | --- | --- | --- | --- | --- | --- | 5424 | 75.99 | 5901564 | 56.41 | Partial | | 86068 |

| | | | | | | | | | | | | | |
|---|---|---|---|---|---|---|---|---|---|---|---|---|---|
| Sesuvium_sesuvioides_gen_lmu | --- | --- | --- | --- | --- | --- | 6689 | 93.71 | 9779056 | 93.48 | --- | --- | --- |
| Sesuvium_verrucosum_trp | --- | --- | --- | --- | --- | --- | 6049 | 84.74 | 8265334 | 79.01 | Partial | | 92783 |
| Spinacia_oleracea_gen | --- | --- | --- | --- | --- | --- | 6457 | 90.46 | 8663105 | 82.81 | --- | --- | --- |
| Stegnosperma_halimifolium_trp | --- | --- | --- | --- | --- | --- | 5270 | 73.83 | 6856989 | 65.55 | --- | --- | --- |
| Trianthema_portulacastrum_trp | --- | --- | --- | --- | --- | --- | 6247 | 87.52 | 6672024 | 63.78 | Partial | | 74484 |
| Trianthema_portulacastrum_trp2 | --- | --- | --- | --- | --- | --- | 6633 | 92.93 | 8140220 | 77.81 | Partial | | 73356 |
| Zaleya_pentandra_trp | --- | --- | --- | --- | --- | --- | 5996 | 84 | 6701387 | 64.06 | Partial | | 62659 |
